# Geographic migration and vaccine-induced fitness changes of *Streptococcus pneumoniae*

**DOI:** 10.1101/2023.01.18.524577

**Authors:** Sophie Belman, Noémie Lefrancq, Susan Nzenze, Sarah Downs, Mignon du Plessis, Stephanie Lo, The Global Pneumococcal Sequencing Consortium, Lesley McGee, Shabir A. Madhi, Anne von Gottberg, Stephen D. Bentley, Henrik Salje

## Abstract

*Streptococcus pneumoniae* is a leading cause of pneumonia and meningitis worldwide. Many different serotypes co-circulate endemically in any one location. The extent and mechanisms of spread, and vaccine-driven changes in fitness and antimicrobial resistance (AMR), remain largely unquantified. Using geolocated genome sequences from South Africa (N=6910, 2000-2014) we developed models to reconstruct spread, pairing detailed human mobility data and genomic data. Separately we estimated the population level changes in fitness of strains that are (vaccine type, VT) and are not (non-vaccine type, NVT) included in the vaccine, first implemented in 2009, as well as differences in strain fitness between those that are and are not resistant to penicillin. We estimated that pneumococci only become homogenously mixed across South Africa after about 50 years of transmission, with the slow spread driven by the focal nature of human mobility. Further, in the years following vaccine implementation the relative fitness of NVT compared to VT strains increased (RR: 1.29 [95% CI 1.20-1.37]) – with an increasing proportion of these NVT strains becoming penicillin resistant. Our findings point to highly entrenched, slow transmission and indicate that initial vaccine-linked decreases in AMR may be transient.

**One-Sentence Summary:** We describe geographic migration, and fitness dynamics conferred by NVT strains and AMR, for the globally endemic pathogen *Streptococcus pneumoniae*.

---

The greatest public health burden from infectious diseases remain stubbornly endemic pathogens. Once established, diseases such as tuberculosis, HIV, and now COVID-19, are very difficult to control, even when vaccines are available(*1*). Their persistence in the population can be partially explained by co-circulation of multiple strains of the same pathogen. This complicates the study of these persistent endemic pathogens in that we rarely understand the mechanisms driving spread, including the role of human behaviour, or why some lineages increase in prevalence over time, while others disappear. Underlying genetic diversity is particularly extreme in the case of the bacterium *Streptococcus pneumoniae* (the pneumococcus), which is the leading cause of morbidity and mortality due to lower respiratory infections worldwide(*2*). The pneumococcus comprises >100 known antigenically distinct serotypes and over 900 classified lineages (also known as global pneumococcal sequence clusters; GPSCs)(*3*); it is not uncommon for over 30 antigenically distinct serotypes to cocirculate within a country or region, or for a human host to carry multiple serotypes concurrently(*4*). Here we develop mathematical models, applied to thousands of geolocated genome sequences from South Africa, collected over a 15-year period to unlock several key uncertainties in pneumococcal migration including the rate and breadth of mobility geographically, and how fitness changes linked to vaccine implementation and antimicrobial resistance may impact its spread.

The pneumococcus resides in the human upper respiratory tract. Carriage is a prerequisite for disease and rates of carriage in children under 5 years-old range from 20-90%(*5*). Occasionally, asymptomatic carriage goes on to cause local infections such as otitis media, or more severely, invasive pneumonia and meningitis. Approximately 300,000 annual deaths linked to pneumococcus are estimated to occur globally(*6*). Penicillin was first used to treat pneumococcal disease in the 1930s and successfully reduced pneumococcal disease until the late 1960s when penicillin non-susceptibility was first noticed. Multi-drug resistant (MDR) strains were described soon after penicillin resistance(*7, 8*). By 2019, 19% of deaths associated with antimicrobial resistance (AMR) had pneumococcal aetiology(*9*). In this context, vaccines are pivotal to disease control. Pneumococcal conjugate vaccines (PCVs) target a small subset of the polysaccharide capsular serotypes, with the most common formulations having included PCV7 and PCV13 (Pfizer)(*10*) and PCV10 (GSK)(*11*). These target 7, 13, and 10 serotypes respectively (with all the serotypes included within PCV7 and PCV10 also included within PCV13). The vaccine serotypes were selected due to their high prevalence and antimicrobial resistance among disease isolates from infants and children. PCVs are now included in 76% of countries’ national immunisation schedules with different formulations in different countries. For example, in South Africa PCV7 was implemented in 2009 and replaced by PCV13 in 2011, excluding PCV10 (*12*). Despite their success at reducing disease, their use has been linked to serotype replacement by non-vaccine serotypes (NVTs) in both IPD and carriage (*4*). In South Africa this has been characterized by increases in NVT serotypes 8 and 15B among IPD, and increases in NVT serotypes 16F, 24, 35B, and 11A among carriage isolates(*13, 14*). At the population level, quantitative measures of serotype-specific fitness changes following vaccine implementation are lacking. This includes quantifying the time it takes after implementation for vaccines to impact the serotype composition in the country as well as quantifying the serotype changes pre- and post-vaccine at different time points. In addition, vaccine implementation has resulted in reductions in antimicrobial resistance among both IPD and carriage isolates (*13–15*). It still, however, remains unclear if these reductions will persist over time at the population level, or if AMR may rebound.

Mathematical models applied to geolocated pathogen genome sequence data provide a powerful tool to disentangle the changing prevalence of different lineages. However, traditional phylogeographic models focus on the rate of pathogen flow between places, and cannot be easily linked to individual behaviour of infected people and the surrounding population(*16*). Additionally, they are unable to account for changing levels of surveillance in both space and time; a common issue in genetic analyses. We have developed models that use the generation time distribution to translate time-resolved phylogeny branch lengths into individual transmission events. Together with measures of human mobility probabilities and human population distribution we can elucidate mechanisms of pneumococcal migration. We implemented this model with a focus on South Africa where approximately 65% of children ≤5 years of age (35% across all age groups) carry the pneumococcus(*17, 18*). We incorporate both the phylogeny and sampling uncertainty through a bootstrapping approach(*19*).

## Results and Discussion

### Quantifying Pneumococcal Spatial Structure

In partnership with the South African National Institute for Communicable Disease (NICD), and the Wits Vaccines and Infectious Diseases Analytics Research Unit (Wits-VIDA), we sequenced the whole genomes of isolates from each of South Africa’s nine provinces between 2000 and 2014 (N=6910, 5060 from individuals with invasive pneumococcal disease and 1850 from carriage studies)(Figure 1A-C) from which we identified 184 GPSCs with 69 different serotypes (31.9% NVT)(Figure 1A, Figure 1E, Table S1). This diversity persisted across provinces and the distribution of serotypes within GPSCs did not follow a distinct geographic structure (Figure 1A). Antimicrobial resistance predicted in-silico was common, (penicillin: 48.2%, erythromycin: 17.3%, clindamycin: 11.2%, and co-trimoxazole: 68.4%) with similar distributions for isolates from both carriage and disease (Figure 1F, Figures S15-S16, Table S3).

**Figure 1.**
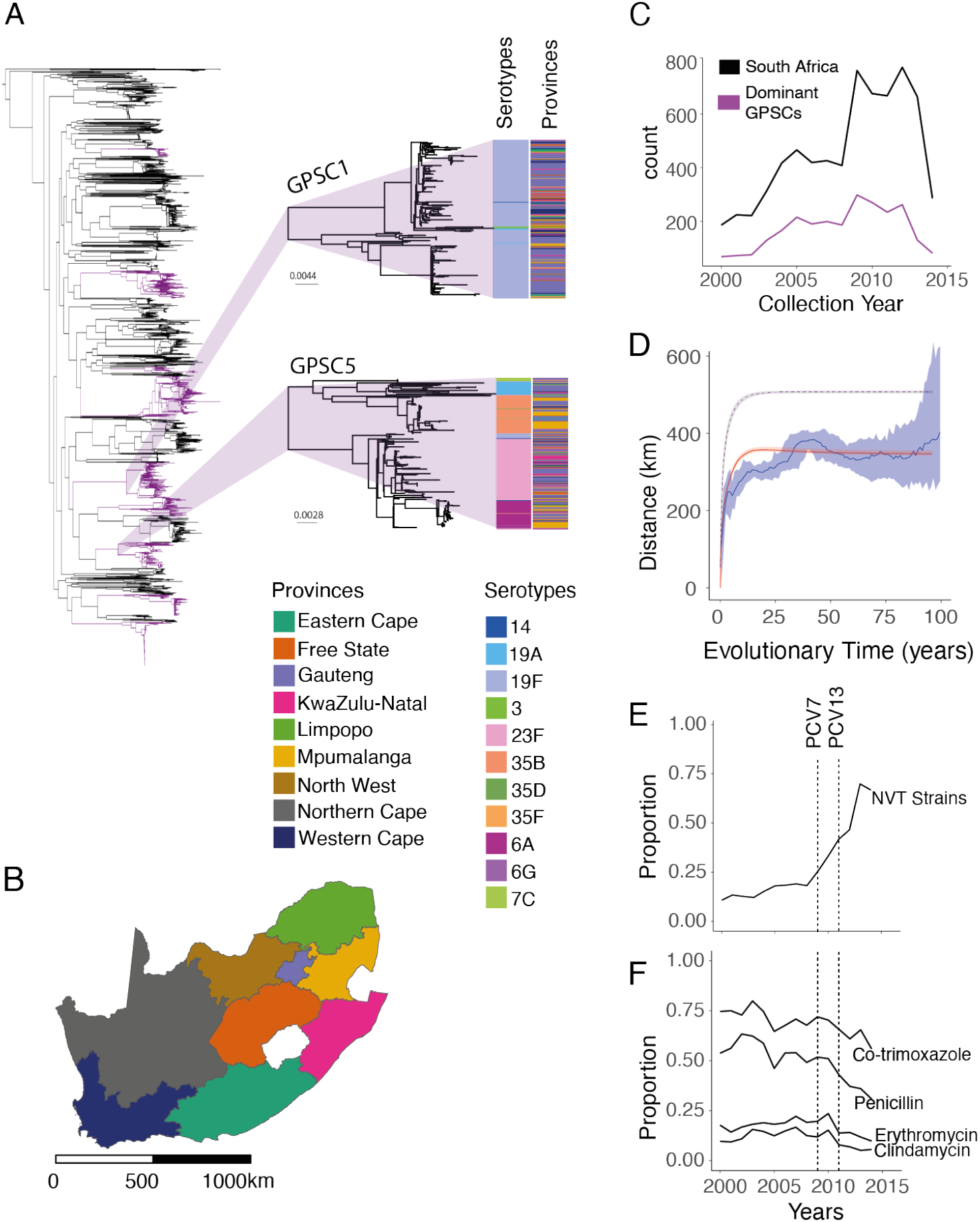
**(A)** Phylogenetic tree of 6910 South African isolates included in this study. Dominant GPSCs (N>50) are coloured purple. GPSC1 (top) and GPSC5 (bottom) are highlighted. The columns describe the serotypes and provincial region for each isolate respectively. **B)** Map of the 9 provinces of South Africa coloured by provinces. **C)** Count of isolates (N=6910) per collection year from 2000-2014 used in the lineage level analysis (black), and the 9 Dominant GPSCs used in the divergence time analysis (maroon). **D)** The mean geographic distance for sequence-pairs as a function of cumulative evolutionary distance across all GPSCs with 95% confidence intervals (blue). The model fit is shown in red. The implied true pattern of spread is shown in purple, after accounting for biased observation process. **E)** The proportion of non-vaccine type serotypes across the study period. **F)** The proportion of antimicrobial resistant isolates for 4 drugs across the study period. The vertical lines denote the introduction of PCV7 in 2009 and PCV13 in 2011. An interactive phylogeny and metadata are available on Microreact. (**https://microreact.org/project/4g3tQZkt4ncWWV9ybPaG3W-southafrica6910)**

Taking the nine most dominant GPSCs in turn (each comprising more than 50 sequences, 2575 sequences total), we built recombination-free, time-resolved phylogenetic trees to determine the divergence times between sequence pairs (Figure S1A-I, Table S4). We included 1157 genomes from 14 other countries in Africa and 2944 from 31 countries outside Africa in our phylogenies. We compared the geographic distance spread per divergence time between pairs in South Africa and found a clear geographic structure with the geographic distance between pairs rising from a mean distance of 142km [95% CI 54-207km] for those separated by less than 2 years of evolutionary time to 297km [95% CI 274-323km] for those separated by 10-20 years (Figure 1D). We obtained consistent results when using only sequences that came from individuals suffering disease (Figure S2A, Table S5), and across the different GPSCs (Figure S3). This is consistent with largely common patterns of spread irrespective of which particular GPSC an individual is infected with. Despite high heterogeneity in GPSC composition within any province, overall, pairs of isolates which are from the same province had 1.27 [95%CI 1.23-1.31] times the relative-risk (RR) of being the same GPSC compared with pairs of isolates from distant provinces (>1000 km). This relative risk fell to 1.09 [95%CI 1.03-1.12] for pairs separated by 500-1000km (Figure 2A, Table S5). We got consistent results when we limited our analysis to only disease isolates and when we subsampled to have even numbers of sequences by province to mitigate sampling bias (Figure S4A-J, Table S5). As there can be hundreds of years of diversity within a single GPSC, we refined the analysis by using different evolutionary windows of separation between pairs of isolates, as determined from the phylogenetic trees. We found that pairs from the same province had 2.43 [95%CI 2.02-3.17] times the RR of having a recent common ancestor (within 5 years) than distal pairs (>1000km apart) (Figure 2B). As the evolutionary time between isolates increases the relative probability of being from the same province decreases, however, it is only after around 50 years that the pneumococcus appeared to be well mixed throughout the country (Figure 2C-E, Figure S5A, Table S6). Further, there appears to be limited flow outside the country, either to other African countries, or to non-African countries, as pairs of isolates separated by 50-200 years had a higher risk of both being from distal South African provinces than pairs from South Africa as a whole and other countries (Figure 2C-E, Figure S5B, Table S5). Recognizing that our samples span age groups we stratified the analysis by the age difference between pairs and found that genome pairs from individuals who were greater than 5 years apart in age took slightly longer to become mixed across South Africa (Figure S5C). These findings are consistent with a highly entrenched pathogen, that moves slowly within a country and with slow cross-border transmission.

**Figure 2.**
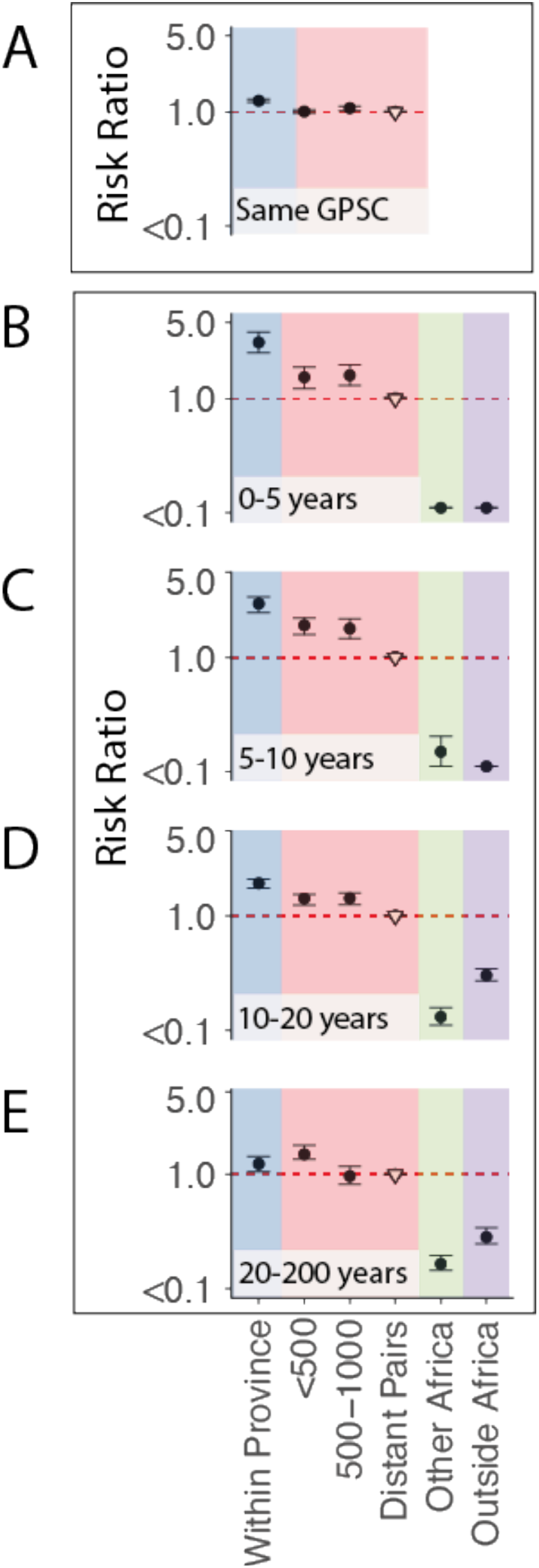
Risk ratio framework to determine geographic structure. (A) Risk ratio of being the same GPSC within a province (blue), between different provinces over increasing distance (red), compared to geographically distant pairs (>1000km) (reference). (South Africa; N=6910). **(B)** Risk ratio of having tMRCA 0-5y, **(C)** 5-10y, **(D)** 10-20y, and **(E)** 20-200 years ago within South African provinces (blue), across larger distances within South Africa (red), from South Africa to other countries in Africa (N=1157) (green), and from South Africa to countries outside of Africa (N=2944) (purple). All plots use a reference of pairs which are from distant provinces in South Africa (open triangle).

### Inferring Mechanisms of Migration using Human Mobility

To understand whether human mobility can explain the slow spread of the pneumococcus, we built a mechanistic model of geographic spread, fit to the observed province in which our genome sequences were isolated. We used the generation time distribution, time from one person being infected to infecting the next person (estimated mean of 35 days, standard deviation of 35 days [gamma distribution]) to translate branch lengths to the number of generations between pairs of sequences (Figure S6)(*20, 21*). Each transmission generation is a possible transmission event and an opportunity for pneumococcal mobility. We use directional human mobility probabilities between each of the 234 South African municipalities from Meta Data for Good (*19*) to infer the probable location of a single transmission event, allowing for mobility of both the infected individual and the surrounding susceptible population. As Meta users may act differently to those involved in pneumococcal transmission, we incorporated a parameter that allows individuals to have a different probability of staying within their home municipality than Meta users (Figure 3B). We then calculated the probability of pneumococcal movement between each pair of locations for each transmission generation. This approach integrates over all possible pathways linking two locations. We incorporate the probability of an isolate being sequenced at each geographic location, within each collection year to account for the number of isolates being sequenced differing by year and location. We fit the model in a Bayesian framework using Markov chain Monte Carlo (MCMC).

**Figure 3.**
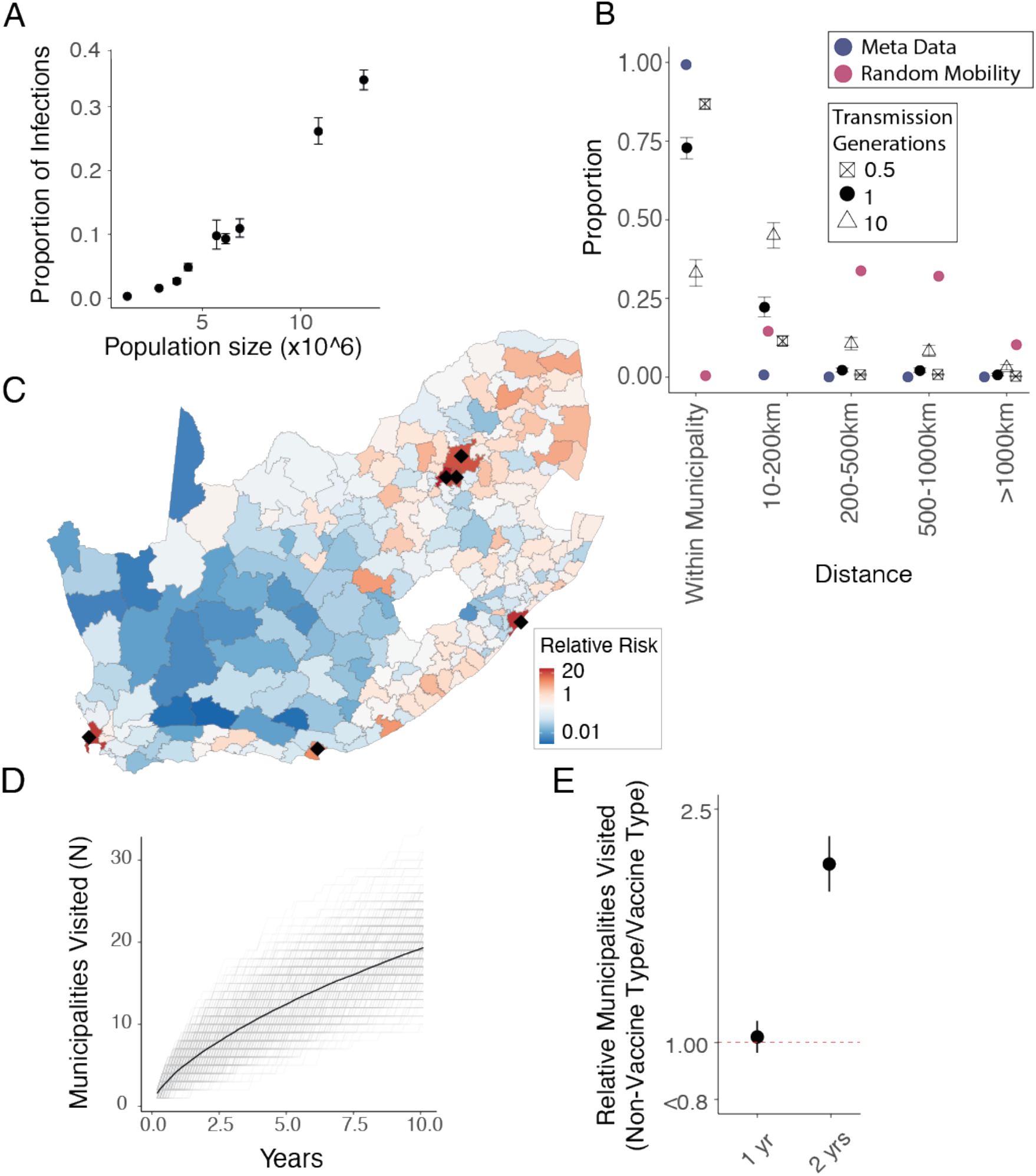
**(A)** The estimated probability of the location (province) of each MRCA (y-axis) compared to the population size (x-axis) in that province. **(B)** The distance from the origin location of a single isolate (0.5 transmission generations). The distance between the original location of a strain and its location after 1 and 10 transmission generations. This is compared to distances if mobility was uniformly random (red) as well as the unadjusted Meta data mobility (blue). **(C)** The relative risk of being in each of the 234 municipalities of South Africa after 1 year (10 transmission generations) of sequential person to person transmission as compared to being in a randomly selected municipality. Black dots denote municipalities with populations of >3 million people. **(D)** The number of unique municipalities visited for 500 unique sequential simulations (grey) and the mean (black-dashed) across years of transmission following an introduction in a randomly selected municipality given the modelled migration probabilities at each transmission generation. **(E)** The relative number of unique municipalities visited by NVT serotypes compared to VT serotypes after 1 and 2 years of transmission.

Our model was able to recover the spatial spread in pneumococcus with increasing evolutionary separation, given the observed number of isolates obtained by location and year (Figure 1D; red line). Our models parameter estimates showed that the relative proportion of the population carrying the pneumococcus per province at any time was strongly correlated with population size in that province (R^2^=0.97; p=<0.01)(Figure 3A). This model allows us to infer the true underlying spread of pneumococcus (i.e., in which we account for the biased observation process) (Figure 1D; dashed line). We estimate that among individuals involved in pneumococcal transmission, the probability of staying in their home municipality was 87.6% [95%CI 80.2-88.7%](Figure 3B; black squares) as compared to 98.9% (ranging from 86.6% in Mogale City, Gauteng to 99.9% in Ba-Phalaborwa, Limpopo) for Meta users (Figure 3B; blue points). When we incorporate the mobility of both infector and infectee, we estimate that after a single transmission generation, 72.8% [95% CI 69.4-75.7] of strains remain in their starting municipality, 22.2% [95%CI 19.3-25.6%] are in a neighbouring municipality, and a small minority are more than 500km away (Figure 3B; black points). As the number of transmission generation increases the probability of reaching distal municipalities increases (Figure 3B). The size of the community appears key to determining where lineages travel. After one year of sequential transmission, we find the probability of being in a municipality with a population size >3 million people was 25.1 [95%CI 18.0-35.2] times that of being in a randomly selected municipality, consistent with a majority of pathogen movement passing through urban centres (Figure 3C, Table S7).

The municipality in which a strain emerges also appears important. After one year of sequential transmission a new strain which first occurred in a rural municipality (population density <50 people/km^2^) has travelled a median distance of 472.7 [95%CIs 74.9-1240.5] kilometres, whereas in the same time window a variant first occurring in an urban municipality (>500 people/km^2^) has travelled only 277.9 [95% CIs 36.3-916.9] kilometres. Further, the variant which emerged in a rural municipality would have travelled to 1.45 times as many municipalities as the urban variant (Figure S7A-C). This is corroborated by previous research which demonstrated high levels of in and out migration amongst individuals in rural settings due to travel for work or education (*22, 23*). On average, one year after emerging, transmission chains have visited 4.34 [95% CIs 2-7] municipalities, and after ten years they have visited 19.14 [95% CIs 12-27] municipalities (Figure 3D). Overall, we find the breadth of geographic spread is driven by a small number of long-range transmission events with most transmissions remaining local. Incorporating our model into a branching epidemic we find that after 10 years; transmission events are an average 398km (366-431) from where they began (Figure S8A-B). We used a simulation approach to test the performance of our model. We simulated spread with known parameters and biased observation processes. We then fit our model on the resulting dataset and were able to recover the underlying true geographic spread between locations (Figure S9).

### Fitness Impact of Vaccine and Antimicrobial Resistance

We found that the implementation of PCV7 in 2009 and PCV13 in 2011 was associated with a substantial disruption in the patterns of circulating serotypes, consistent with what has been observed elsewhere (*4, 13, 24, 25*). By 2014, serotypes included in PCV13 represented 33.2% of all isolates in our dataset, down from 85.0% in the pre-vaccine era (Figure 4A). These patterns were very consistent across the nine provinces in South Africa (Figure S10). To quantify changes in fitness linked to the vaccines, we fitted models to the annual distribution of serotypes, allowing for differential fitness in serotypes included in PCV7 (serotypes: 4,6B,9V,14,19F,18C,23F), PCV13 (additional serotypes: 1,3,5,6A,7F,19A), and those not included in the vaccine (NVT). This method tracks the proportion of all serotypes at the population level over time, and quantifies the relative advantage of each of them following the implementation of the vaccine. This simple formulation was able to recover the observed distribution of serotype proportions in each group, by year, across provinces (Figure 4A-C, Figure S10). We note that the number of NVT and VT isolates we used in our model do not represent the underlying incidence of NVT and VT, as only a small proportion of all infections were detected and sequenced. However, as we focus on the relative abundance of NVT and VT strains per year, our approach is robust to changes in the absolute numbers of isolates sequenced.

**Figure 4.**
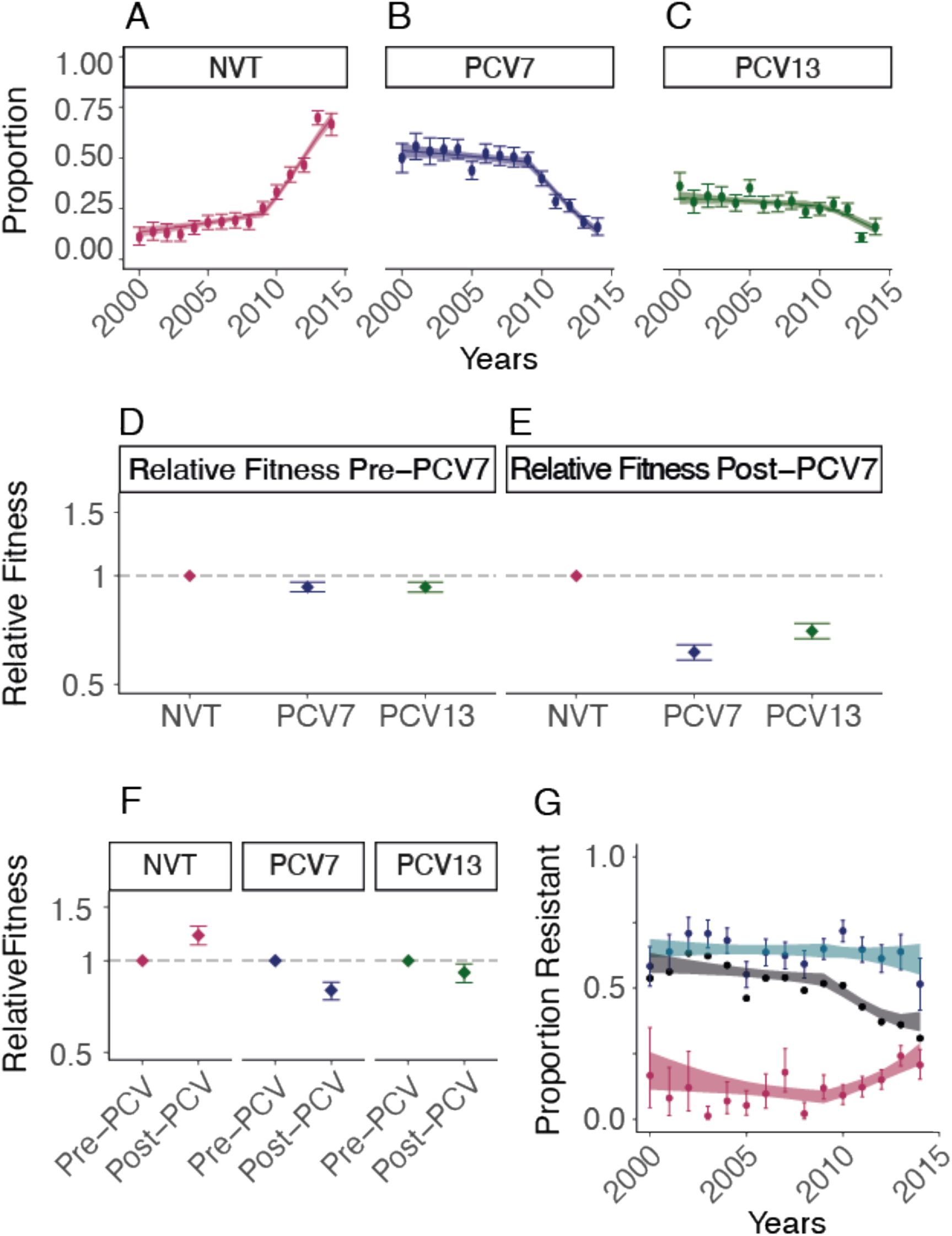
Pneumococcal fitness estimates. Data (points) and model fit (line) for the proportion of serotypes from **(A)** non-vaccine types, **(B)** PCV7 types, and **(C)** additional PCV13 types not included in PCV7 from the years 2000 to 2014 in this study. **(D-E)** Relative fitness for the three groups of serotypes compared to the NVT fitness estimates in **(D)** the pre-PCV era (pre-2009) and **(E)** the post-PCV introduction era (post-2009). **(F)** Relative fitness estimates for all three groups of serotypes comparing the post-vaccination (post-2009) to the pre-vaccination (pre-2009) eras. **(G)** Proportion of penicillin resistance overall (black line), within NVT strains (maroon points), and within VT strains (turquoise points) with model fits.

We found that prior to the implementation of the vaccine (2000-2008), NVTs had a relative fitness of 1.06 [95% CI 1.04-1.07] when compared to serotypes included in the vaccine (combining serotypes in PCV7 and PCV13 into VT). We found that the introduction of the vaccine is associated with a marked change in fitness. When comparing serotype fitness pre- and post-introduction of PCV, VTs had a relative fitness 0.80 [95% CI 0.74-0.85] and 0.91 [95% CI 0.85-0.98] for PCV7 and PCV13 serotypes respectively. Meanwhile, among NVT serotypes the vaccine was associated with a 1.28 [95% CI 1.21-1.30] times increase in relative fitness from 2010-2014, as compared with prior to the vaccine (Figure 4D-E). The fitness of VTs relative to NVTs declined from 0.93 [95% CI 0.90-0.96] in the pre-PCV era to 0.66 [95% CI 0.63-0.69] after vaccine implementation (2009) (Figure 4F, Table S8). When we directly compare the fitness advantage of NVTs compared to VTs in the PCV era, NVTs have a relative fitness advantage of 1.53 [95% CI 1.47-1.60], which is equivalent to 1.04 [95% CI 1.04-1.05] growth advantage (relative to VTs) at each transmission generation. In a sensitivity analysis, we show that the results remain consistent for carriage and disease isolates respectively, despite them having been sampled from different cohorts (Figure S11). We also find consistent results across provinces (Figure S10).

We were able to refine this model to look at the fitness of individual serotypes and found a wide range of fitness across serotypes. Among NVTs serotypes 15A, 35B, and 8 had the greatest fitness advantage post-PCV (Figure S12-14) concordant with what has been previously described (*13, 14, 26*). We also found that shifts in lineage fitness result in changing patterns of spread. By incorporating our fitness estimates for NVT and VT strains after vaccination into our mobility model we estimated that the number of affected municipalities from a strain of a non-vaccine type was 2.02 [95%CI 1.81-2.25] times the number of affected municipalities from types included in the vaccine (Figure 3E, Figure S8C-F).

The serotypes included in the vaccines were prevalent in childhood disease and had high levels of AMR in the United States, where the vaccines were developed (*27*).The high levels of AMR in the VT strains was also present globally(*15*). In South Africa, similar to other countries, reduction of AMR has been noted since vaccine implementation; however, it remains unclear whether this trend persists or if AMR eventually rebounds(*15*). In South Africa, prior to the vaccine, 63.6% of VT and 8.8% of NVT strains were predicted to be penicillin resistant. We found there was a clear drop in overall penicillin resistance following vaccine implementation, driven by reductions in the proportions of strains that are VT (Figure 4G, Table S9). The trends, while still present, were less clear in the other investigated antimicrobials. Using our same modelling framework, we were able to recover the observed proportions of strains that were resistant to penicillin over time. We found that prior to the vaccine, among both NVT and VT strains, there was limited difference in the fitness between those that were penicillin-resistant and penicillin-susceptible. However, following implementation of the vaccine, NVT resistant strains were 1.30[95% CI 1.19-1.43] times as fit as penicillin-susceptible NVT strains (Figure S15A-D). Contrarily resistance did not appear to have changed among VT penicillin-resistant strains (relative fitness of 0.97 [95% CI 0.91-1.03])(Figure 4G, Figure S16A-C, Table S9). Expansion of NVTs within typically VT-associated lineages is the most common mechanism for serotype replacement(*25*). As a result, the penicillin resistance associated with these newly expanded NVT lineages is able to persist in the population(22). Together with our findings, this suggests that the overall reduction in penicillin resistance seen following vaccine implementation may not persist. This highlights the nuanced effect that vaccines can have on patterns of AMR (Figure S16D-F). It is likely that next generation higher valency PCVs may also lead to increased AMR prevalence in those serotypes not included.

## Limitations

We did not have complete carriage and invasive disease data across the entire time period. In addition, the human mobility data did not overlap with the sequence collection dates. By adjusting the probability of being at home in the human mobility data we address the possibility that the Meta mobility data does not reflect the true movement of individuals carrying pneumococcus in these years. We also performed sensitivity analyses to determine whether our results are robust to both carriage and invasive disease. Within the fitness model, as we are looking at relative proportions of strains with increasing proportions of resistance, there may still be decreased total burden of resistant disease if those strains carrying it remain low prevalence. We are unable to address this with the data we currently have.

## Conclusion

Here we describe the movement of a persistent human pathogen whose geographic course has been largely hidden by its diversity and endemicity. The pneumococcus has an affinity for urban centres through which it channels its wider geographic spread. Although it is characterized by slow transmission overall, the perturbation of vaccines can substantially and rapidly change pneumococcal lineage ecology. Although vaccine-associated fitness dynamics have been previously identified in the pneumococcus they have not been directly quantified (3,15). Increasing proportions of NVTs in the PCV era invasive disease isolates can be largely attributed to the decrease in number of VTs rather than increasing prevalence of NVTs(*14*). Vaccination has had a secondary effect on penicillin resistance with a decrease in recent years in South Africa. Given our estimated growth advantage of penicillin-resistant NVT strains we may see a reversal of this benefit. Further, we have quantified pneumococcal geographic spread and the spatial impact of NVT expansion after vaccination. This mechanistic description of pneumococcal geographic spread provides insight into the movement of emergent strains. In South Africa, the population density in the emergence location of a NVT strain may impact its speed of spread across the country and have implications for public health responses to emergent strains. Emergence in a highly mobile, peri-urban area may enable both rapid proximal geographic spread and less frequent distal seeding events, and thus more complete distribution across the country. The fitness model and human mobility model together provide frameworks to quantify and better understand the migratory and fitness dynamics of this globally endemic pathogen. The magnitude of South Africa and its provinces demonstrates that these frameworks may be applied to other large regions.

## Supporting information

Table S1 will be used for the link to the file on the preprint site.

## Acknowledgments

This work was supported by Wellcome under grant reference 206194 and by the Bill and Melinda Gates Foundation under Investment ID INV-003570), NIH (HS, grant number R01AI160780) and the European Research Council (HS, NL, grant number 804744). For the purpose of Open Access, the author has applied a CC BY public copyright licence to any Author Accepted Manuscript version arising from this submission

Further we would like to acknowledge Megan O’Driscoll, Dr. Stephen Farr, and Dr. Victoria Carr for code review.

## Author contributions

Conceptualization: SB, HS, SDB

Methodology: SB, NL, HS, SDB

Visualization: SB, NL, SDB, HS

Sample Collection & Processing: SN, SD, AVG, SAM, LM, MDP

Funding acquisition: SB, SDB, HS

Supervision: SDB, HS

Writing – original draft: SB, SDB, HS

Writing – review & editing: SB, NL, SN, SD, MDP, SL, GPS, LM, SAM, AVG, SB, HS, AP, CMA, KK, JL, NC, DL, RK, RD, SM, BV

## Data and materials availability

All data are available in the main text or the supplementary materials. All scripts for analysis are available on GitHub at: https://github.com/sophbel/Analysis_GeographicMobility_pneumoPaper

