## Supplementary material for "Geographic migration and vaccine-induced fitness changes of *Streptococcus pneumoniae*": Table S1 will be used for the link to the file on the preprint site.

**This PDF file includes:**

Materials and Methods: 3-18
Figs. S1 to S24: 19-43
Tables S1 to S9: 44-54

### Materials and Methods

### ***Data Sources & Processing***

#### Pneumococcal sequence data and metadata

The genomes included in this study were collected as part of the Global Pneumococcal Sequencing project (GPS); a global genomic survey of *Streptococcus pneumoniae*(*29*). The invasive disease isolates included here were collected by the National Institute for Communicable Disease (NICD) in South Africa from 2000-2014(*14*). In the initial phase of genome sequencing approximately 300 invasive-disease isolates from each year (2005-2014) were selected with a specific target age breakdown (50% from <3-year-olds, 25% from 3-5-year-olds, 25% from >5-year-olds). In the second phase of sequencing approximately 200 disease isolates from 2000-2004 and approximately 100 invasive disease isolates from 2005-2010 from children <5 years old were randomly selected for sequencing(*14*)(Table S2). The carriage isolates were collected in Soweto, Gauteng (N=736; collection years: 2010, 2012, 2013) and Agincourt, Mpumalanga (N=1114; collection years: 2009, 2011, 2013) by the Wits Vaccines and Infectious Diseases Analytics Research Unit (Wits-VIDA). A random sample of 400 carriage isolates from each year were chosen for genome sequencing (Table S2). We included both carriage and invasive disease isolates and conducted sensitivity analyses throughout to confirm the methods were robust to both carriage and invasive disease individually. Invasive disease is defined as the bacterium being isolated from a typically sterile site. The majority of total isolates were from children aged <5 years (80.5%), while 13.8% were from children aged 5-18 years and only 5.7% were from adults (>18 years). These were distributed across the pre- and post-PCV periods. We additionally included similar GPSCs from the GPS database (for context within the global population) for the relative risk analysis. We utilized metadata including collection year and month, residence province of the patient, age of the patient, sampling site, and clinical manifestation. The range of sampling sites include nasopharyngeal swabs (for carriage); blood, pleural fluid, cerebrospinal fluid, peritoneal fluid, pus, and other joint fluid (for invasive disease).

#### Population Data

We estimated the population for each municipality (N=234) across South Africa using the populations-size estimates from LandScan 2017(*30*, *31*).

### Mobility Data

The mobility data used in this study was collected using Meta Data for Good Disaster maps from South Africa. These are initiated at the onset of a disaster – in this case the SARS-CoV2 pandemic and track the geographic movement of Meta users(*32*). We used the baseline (adjusting to 2 weeks prior) human mobility for each month from January 2020 to July 2021(*32*, *33*) to attain a mobility probability from and to each municipality. For each origin (home) municipality (N=234), we determined the mean monthly number of Meta users that were in each destination municipality in South Africa. Location pairs with a value of zero were given a value of 10. We divided each cell by the total number of users across all destinations for that origin municipality. Each cell in the resultant original-destination matrix therefore represented the probability of being in each destination municipality, given your home municipality. This is the probability of mobility from each municipality to each other municipality after 2019. Because we don't have mobility data from the exact years the genomes were sampled we adjust the diagonal of the mobility matrix using an estimated parameter. This allows people to stay more or less at home and mitigates the impact of non-year matched mobility data.

### Generation Time Distribution

We used a simulation framework to estimate the overall generation time using the separate contributions of the carriage durations and the incubation period. This approach has previously been employed for other pathogens(*34*).

We sample 1000 carriage durations from an exponential distribution with means which are inverse to the clearance rates estimated in Chaguza, C., et al. 2021(*20*) (Clearance Rate: 0.026 [95% CI: 0.025–0.028 episodes/day), and Abdullahi, O., et al., 2012 (Clearance Rate: 0.032 episodes/day [95% CI: 0.030–0.034])(*21*).

To sample the day of transmission, we then randomly sample a time point between 0 and the time of clearance for each individual. We then separately sampled an incubation period using a uniform distribution of between 1 and 5 days. To account for longer carriages resulting in more opportunities for transmission, we sampled from the distribution of generation, with the probability of sampling each individual weighted by the total carriage duration. The total generation time is then the sum of the duration to transmission and the incubation period.

We repeat these steps 10,000 times and estimate the mean and standard deviation of this final distribution assuming that the generation time follows a gamma distribution with mean 35 and standard deviation 35 (exponential distribution with a rate of 1/0.096). By comparing the histogram of the distribution with the gamma distribution, this appears a reasonable assumption (See Figure S6).

#### *Sample culture and genome sequencing*

These pneumococcal isolates were selectively cultured on BD Trypticase Soy Agar II with 5% sheep blood (Beckton Dickinson, Heidelberg, Germany) and incubated overnight at 37°C in 5% CO2. Genomic DNA was then extracted manually using a modified QIAamp DNA Mini Kit (QIAGEN, Inc., Valencia, CA) protocol. As part of GPS, pneumococcal isolates were whole-genome sequenced on the Illumina HiSeq platform to produce paired-end reads with an average of 100-125 bases in length and data were deposited in the European Nucleotide Database. Whole genome sequence data was processed as previously described(*25*).

#### *Antimicrobial resistance*

We performed predictive antimicrobial susceptibility profiling using the CDC-AMR pipeline for 3 classes of antimicrobials: beta-lactams (penicillin; encoded by the genes pbp1A, pbp2B, pbp2X)(*35*, *36*), sulphonamides (co-trimoxazole; *folA* and *folP*), macrolides (erythromycin and clindamycin; *ermB* and *mefA*)(*37*, *38*). This was done for 6,798 randomly selected isolates(*29*).

#### *Constructing time resolved phylogenetic trees*

We selected the GPSCs for which we had genomes from each of South Africa's nine provinces and for which we had a minimum of 50 sequences in total to build phylogenies, henceforth referred to as "Dominant GPSCs". There were nine dominant GPSCs. We created reference genomes for each GPSC using ABACAS to order the contigs from a representative of each GPSC mapped to *Streptococcus pneumoniae* (strain ATCC 700669/Spain 23F-1) [EMBL accession: FM211187]. Any contigs which did not align were concatenated to the end. We multiply mapped all genomes from each Dominant GPSC against these references respectively using a custom mapping, variant calling, and local realignment around indels pipeline using bwa-MEM(*39*) and samtools mpileup(*40*). We built trees masking recombination regions using Gubbins(*41*) with RAxML(*42*) and a GTR model. We converted branch length to time using BactDating with a mixed gamma, relaxed clock model(*43*). We compared concordance between BEAST(*44*) with both strict and relaxed clocks, and a Bayesian skyline prior. As the results were concordant, we used BactDating due to its shorter runtime (Figure S17).

### ***Risk Ratio Framework***

We used a risk ratio framework to investigate the risk of genetic similarity across geographic distance(***45***). We compare the location (*loc*) and label (*G*) (i.e., GPSC or genetic similarity) of pairs of sequences that were collected around the same time (*t*). This approach has been shown to be robust to substantial biases in timing and location of isolate collection(***45***). We first constructed pairwise matrices comparing every isolate to every other isolate (N Pairs=6910). In the numerator was the ratio of pairs which were the same GPSC, collected within a year of each other, from the same province, over the total number of pairs collected within a year of each other from the same province. The denominator was the ratio of pairs which were the same GPSC, collected within a year of each other, from distant provinces (>1000km apart)(*Lref*), over the total number of pairs collected within a year of each other from distant provinces. Geographic distances were calculated based on the centroid coordinates of each province. To demonstrate the suitability of using centroid distances, we simulated a spatial transmission process for 1000 separate chains where at each generation a daughter point is placed at a randomly located location 350m in each of the x and y direction. This is repeated over 20 generations. We then identify the 'centroid' of each case based on the closest coordinate rounded to the nearest kilometre. We then calculate the total distance covered for both the true distance and the centroid distances and found that the resulting distances are very similar (Figure S18).

To quantify uncertainty, we used a bootstrapping approach where in each bootstrap iteration we randomly sampled with replacement the isolates before recalculating the statistic. We report the 2.5 and 97.5 percentiles from the resulting distribution.

We also repeated the same analysis but used the time-resolved phylogenetic trees to interrogate pairs across increasing divergence times (breaking the GPSCs into higher resolution). For this, rather than matrices designating whether pairs were the same GPSC or different GPSCs, the divergence time between each pair was included. We only included the divergence times between like-GPSCs (Figure 2C-E).

Equation 1.

$$RR_{loc}(g1,g2) = \sum_{i=1}^{n}\sum_{j\neq i}^{n} \frac{(loc_i = loc_j \cap t_{ij} \leq 1\, year \cap G_i = G_j) \Big/ (loc_i = loc_j \cap t_{ij} \leq 1\, year}{I(Lref_n = Lref_n \cap t_n \leq 1\, year) \cap G_i = G_j) \Big/ I(Lref_n = Lref_n \cap t_n \leq 1\, year)}$$

We utilized the framework to compare a range of geographic distances keeping the reference distance to pairs which were >1000km apart (Figure 2).

To identify the divergence time at which pairs had an equal risk of being in the same province as distant provinces (time to homogenization across South Africa) we investigated the divergence time at which there was *no* increased risk of similarity within a province compared against distant provinces (Relative Risk = 1). We stratified distances in South Africa into groups of distances including the 9 within province, 14 pairs which were <500km apart, 16 pairs 500-1000 km apart, and 6 pairs provinces >1000km apart. We repeated the framework across rolling 20-year time windows at 10-year intervals from 0-100 years (*g1,g2*)(Figure S5A). We repeated this for pairs where one was from South Africa and the other was from another country in Africa (Figure S5B).

To quantify uncertainty, we used a bootstrapping approach where in each bootstrap iteration we randomly sampled a tree, and then sampled with replacement the sequences before recalculating the statistic. We report the 2.5 and 97.5 percentiles from the resulting distribution.

We repeated all analysis with sub-sampling to 300 to compensate for biased sampling, and only including pairs isolated from patients with pneumococcal disease (Figure S4).

All statistical analysis was performed in R v3.6.2(*46*).

### ***Probabilistic Mobility Model***

#### Overall Strategy

We extend a mechanistic phylogeographic model previously employed by Salje *et. al*, 2021(*16*) to estimate the mobility of the pneumococcus between pairs of municipalities at each transmission step. To infer the probable path of transmission between sequence pairs we used the divergence time and the generation time distribution to estimate the number of transmission generations between pairs of sequences. Each generation is a possible transmission event and provides an opportunity for a mobility event.

The approach ultimately aims to estimate an origin destination matrix for a transmission step, where each cell represents the probability that the pneumococcus is now in location *j* after one transmission step given it was previously in location *i.* As the phylogenetic trees combined with the generation time provide an estimate of how many transmission steps separate pairs of samples, we can use repeated matrix multiplication to integrate over all possible pathways linking two locations (see below for more details). We incorporate the probability of sampling at each geographic location, within each collection year, by GPSC to account for our observation process.

#### Notation

We follow notation implied by Salje *et. al,* 2021(*16*). A pair of isolates, $C_a$ and $C_b$, with sequences, $Seq_a$ and $Seq_b$ included in a phylogeny are found in locations, $L_a$ and $L_b$, and the samples were taken in the years, $T_a$ and $T_b$. The inferred MRCA between $C_a$ and $C_b$ is time $T_m$ and located in $L_m$. The number of transmission generation from the MRCA to $C_a$ is $G_a$ and to $C_b$ is $G_b$.

#### Model fitting

#### Single Transmission Generation

Considering a single transmission generation, the probability that persons *i* and *j* come into contact with each other given *i* lives in location *a* and *j* lives in location *b* can be written as:

Equation 2.

$$P(person\ i\ and\ j\ come\ into\ contact\ |Li = a, Lj = b) = \sum_{k}^{N} P(Vi = k\ |Li = a) \cdot P(Vj = k|Lj = b) \cdot B_k$$

Where $P(V_i = k \,|L_i = a)$ is the probability that individual *i*, whose home location is in *a*, visits location *k* and $P(V_j = k \,|L_j = b)$ is the probability that individual *j* whose home location is in *b* visits location *k,* and $B_k$ is the location-specific probability of transmission for *k*.

At time $\tau$, one infector in *i* in location *a* is expected to transmit to this number of persons in location *b*:

Equation 3.

$$E(number\ of\ persons\ from\ b\ infected\ at\ time\ \tau | L_i = a) = \sum_{k}^{N} P(V_i = k \,|L_i = a)\ \cdot P(V_j = k \,|L_j = b)\ \cdot B_k \cdot S_{b,\tau,gpsc}$$

Where $S_{b,\tau,gpsc}$ is the number of susceptible people in location *b* at time $\tau$ with some lineage=*gpsc.*

The total number of people infected by the infector is:

Equation 4.

$$E(number\ of\ persons\ from\ all\ locations\ infected\ at\ time\ \tau | Li = a) = \sum_{m}^{N}\sum_{k}^{N} P(V_i = k \,|L_i = a) \cdot P(V_j = k | L_j = m) \cdot B_k \cdot S_{b,\tau,gpsc}$$

Conditional on transmission occurring, the probability that the infectee's isolate is taken in location *b* is:

Equation 5.

$$\delta_{a,b,\tau,gpsc} = P(L_j = b | L_i = a) = \frac{\sum_k^N P(V_i = k \,|L_i = a)\ \cdot P(V_j = k \,|L_j = b)\ \cdot B_k \cdot S_{b,\tau,gpsc}}{\sum_m^N \sum_k^N P(V_i = k \,|L_i = a) \cdot P(V_j = k | L_j = m) \cdot B_k \cdot S_{b,\tau,gpsc}}$$

We then create an NXN transmission matrix, $\Delta_{\tau,gpsc,gen=1}$, with N being the total number of locations, containing the transmission probabilities, asymmetrically, between all pairs of locations at each point in time; $\delta_{a,b,\tau,gpsc}$ is element [a,b] of the matrix..

**Human Mobility Characterisation**

We use Meta mobility data (MetMob), as described in the mobility data section above, to characterise mobility between the 234 municipalities of South Africa. We aggregate these to province level (N=9) to fit the model. Initially we extract a 234X234 matrix which sets out the probability that an individual from municipality *a* visits location *k*; again where *k=any municipality*.

MetMob comes from individuals using Meta however this may not be representative of the amount of time spent at home by those involved in pneumococcus transmission. The mean of the diagonal of the Meta Mobility matrix (234X234) is 0.9893 implying, on average, 98.9% of Meta Users stay in their home municipality, $H_i$. To allow for individuals to spend more or less time at home than represented in the MetMob data we incorporated a parameter to adjust the probability of being at home ($\theta$). We adjust the probability of staying home using a standard logistic function and restrict it with bounds of -0.04 and 0.6 to facilitate exploration of a sensible space. The adjustment allowed by the bounds limits the range of movement $H_i$*-0.6* and $H_i$*+0.04*.

The probability of a person remaining in location *a* therefore becomes:
Equation 6.

$$P(V_i = a|H_i = a) = MetMob[a,a] - (-0.04 + (\frac{(\exp(\theta))}{(1+\exp(\theta))}) \cdot (0.6 + 0.04))$$

Where a value of greater than 1.0 is obtained for a specific municipality, this is replaced by a value of 0.999.

We rescale the probabilities so that the sum of all mobility is equal to 1:
Equation 7.

$$P(V_i = a|H_i = a, k \neq a) = MetMob[a,k]/(1 - P(V_i = a|H_i = a))$$

Where *MetMob[a,k]* considers all mobility probabilities from the Meta Mobility data.

The sum of the movements to South African municipalities is equal to 1 which assumes that the analysis contains all possible movements of both the pneumococcus and people implying no external introductions. The outcome of this is that some mobility may be missed especially around the country borders.

**Probability of the pneumococcus being in each location after G transmission generations**
To determine the probability that location *k* contains the home location after *G* transmission generations, we use matrix multiplication which integrates across all possible pathways connecting two locations.

Equation 8.

$$\Delta_{\tau=T_G,gpsc,gen=G} = \prod_{r=2}^{G} \Delta_{\tau=t_{r-1},gpsc,gen=r-1} \cdot \Delta_{\tau=t_r,gpsc,gen=1}$$

Where $t_r$ is the time of generation $G_r$.

**Probability of observing a pair of cases in two specific locations.**

The probability that $C_A$ has home location $L_A$ and $C_B$ has home location $L_B$ is conditional on the sequences being observed in locations $L_A$ and $L_B$ at times $T_A$ and $T_B$. We assume that the location of two cases, $L_i$ and $L_j$, is dependent on the location of their MRCA, $L_m$, and the number of transmission generations separating them from their MRCA, $G_A$ and $G_B$.

The observations processes across locations are independent of each other and each transmission event is independent of other transmission events. The probability of observing a case at $L_i$ at time $T_A$, is not dependent on the number of generations to, or location of, the MRCA. We consider discretized space of the 9 provinces of South Africa resulting in the following equation:

Equation 9.

$$P\left(L_A, L_B \middle| Obs_{L_B,T_B}, Obs_{L_B,T_B}, Seq_A, Seq_B, T_A, T_B\right) = \frac{\sum_{L_m}\sum_{G_A}\sum_{G_B} P\left(Obs_{L_A,T_A}\right)P\left(Obs_{L_B T_B}\right)P(L_A|L_m, G_A)P(L_B|L_m, G_B)P(L_m)P(G_A, G_B|Seq_A, Seq_B, T_A, T_B)}{\sum_{L_i}\sum_{L_j}\sum_{L_m}\sum_{G_A}\sum_{G_B} P\left(Obs_{L_i,T_A}\right)P\left(Obs_{L_i T_B}\right)P(L_i|L_m, G_A)P\left(L_j \middle| L_m, G_B\right)P(L_m)P(G_A, G_B|Seq_A, Seq_B, T_A, T_B)}$$

**Probability of G generations between MRCA and a sequenced isolate**

Under the previous equation $P(G_A, G_B|Seq_A, Seq_B, T_A, T_B)$, represents the generation time distribution

We can extract the joint probability of $C_A$ and $C_B$ being separated from the MRCA by $G_A$ and $G_B$ transmission generations respectively using the above derived generation time distribution and the time resolved phylogenetic trees.

Assuming the generation time is gamma distributed and all transmission events are independent the sum of the gamma distribution is also gamma distributed. Additional we can extract the evolutionary times $E_A$ and $E_B$, separating $C_A$ and $C_B$ from the MRCA. As described in Salje *et al.* 2021, Equation 14 (*16*) we can estimate the probability of *g* transmission events over many trees allowing us to incorporate phylogenetic including evolutionary parameters from the tree structure.

We determined the probability for the number of generations from MRCA for each isolate for 1 to 1000 generations, using the generation time derived above.

**Location of the MRCA (*$P(L_m)$*)**

We estimate the probability that on average, an MRCA is in each of the nine provinces in South Africa. We estimate parameters for each of 8 provinces setting Western Cape aside as a reference and dividing by the total across all 9 to ensure that the sum of the probabilities is 1. This again assumes no external introductions.

**Calculation of Likelihood**

We can calculate the likelihood using all pairs of available sequenced *S. pneumoniae* as described in Salje et al. 2021(*16*). We accounted for the observation process by incorporating the probability of sampling in each location for isolates belonging to each GPSC annually.

**Likelihood Equation**

We calculate the likelihood using all pairs of sequenced pneumococci as follows:
Equation 10.

$$L \propto \prod_{gpsc=1}^{9} \prod_{i=1}^{n_{gpsc}} \prod_{j \neq i} P(L_i, L_j | Obs_{L_i,T_i}, Obs_{L_j,T_j}, Seq_i, Seq_j, T_i, T_j)$$

Where $n_{gpsc}$ are the number of sequences available from GPSC *gpsc*.

**Hamiltonian Markov Chain Monte Carlo**

We used an MCMC approach to estimate our parameters using the package fmcmc implemented in R (*47*). We estimated 9 parameters:

- a parameter that adjusts the probability of staying in the home municipality as compared to Meta mobility data.
- 8 parameters capturing the relative probability that the MRCA of a pair of individuals was in each of the other eight provinces as compared to the province of Western Cape (the reference).

We only used pairs of sequences that were separated by under 10 years of evolutionary time between them. After 10 years there are limited spatial signals remaining as the bacterium has had many opportunities to move. This approach is to make the model computationally tractable. In a sensitivity analysis we showed that increasing the model to 15 years resulted in similar estimates (Figure S19).

**Model performance**

In order to assess the performance of our model, we used a simulation framework. We simulated 50000 pairs of events, locations, and the location of the MRCA between pairs, where the probability of mobility between each pair of locations was determined by the Meta mobility matrix adjusted by a parameter of known value of -2. We tested our ability to recapture this input parameter. To incorporate the biased observation process, we down-sampled the simulated data based on the by-province sampling probabilities from our true data.

We used the down-sampled data to fit our model with 20000 steps of an MCMC with a jump-step of 0.08 in 3 chains. We were able to recapture the down-sampled data utilising the human mobility framework and the estimated parameter for the probability of staying at home, accounting for the sampling probability per province. We then utilised the same human mobility framework and the estimated parameter, excluding the sampling probability and were able to recapture the complete simulated data including the input parameter (Figure S9).

***Model sensitivity analyses***

Further we estimated our parameters only to include isolates from patients with invasive pneumococcal disease (Figure S2B).

To test the impact of a range of generation times on the parameter estimates we also re-ran our MCMC with generation times of 15, 35 (as included in the model), and 55 days (Figure S20).

We confirmed the chains converged for each of the nine parameters estimated (Figure S21).

#### ***Probability guided transmission simulations***

##### Person-to-person transmission chains

We simulated person-to-person transmission chains seeded in a starting municipality weighted by the population size. We determined the relative risk of being in a specific municipality after 10 transmission generations (approximately one year) across 500000 simulations.

We fixed the starting municipality to be rural (population density <50/km$^2$) or urban (population density >500/km$^2$) and repeated the above simulation. For 10000 sequential simulations we counted the number of unique municipalities affected and distance travelled at each transmission interval weighting by population size. We then determined the number of municipalities travelled to across all transmission chains (Figure S7).

##### Branching epidemic

We simulated a branching epidemic in which we drew the number of transmission events seeded by each event from a Poisson distribution around an effective reproductive number ($R_{eff}$) of 1 and amplitude of 0.15 over 100 iterations. We determined the mean distance from the starting municipality after 60 generations (5.8 years with a 35-day generation time) and the number of municipalities visited over that time. We calculated the uncertainty at the 95% confidence intervals of a normal distribution.

### ***<u>Population-level fitness model</u>***

#### Overall Strategy

We developed logistic growth models to fit the changing prevalence of different serotypes utilizing a method which has previously been implemented for the endemic bacterium, *Bordetella pertussis*(*48*).

#### Vaccine status

We first binned serotypes into three groups those not included in the vaccine (NVTs), serotypes included in PCV7 (4,6B,9V,14,18C,19F,23F), and additional serotypes included in PCV13 (1,3,5,6A,7F,19A). We computed the relative abundance $f_{i,ref}$ of each group of serotypes, *i*., compared to a reference type *ref*.

$$f_{i,ref}(t) = \frac{N_i(t)}{N_i(t) + N_{ref}(t)}$$

We chose to use NVT's as the reference. Varying the reference does not impact the model, as long as the reference samples span all years. We then used a simple logistic model, assuming constant growth rate, to capture the evolution of this abundance, at each time *t*.

Equation 11.

$$f_{i,ref}(t) = \frac{1}{1 + \left(\frac{1 - f_{i,ref,0}}{f_{i,ref,0}}\right) \exp(-r_{i,ref} \cdot t)}$$

where $r_{i,ref}$ is the growth rate of that abundance, shared across all provinces and $f_{i,ref,0}$ is the initial relative abundance of the serotype group $i$ with respect to a chosen $ref$.

To control for the varying presence of all circulating serotypes through time, we present fitness as the average relative growth rate, $\overline{r_i}$, for each group with respect to a randomly selected group in the population:

Equation 12.

$$\overline{r_i} = \sum_{j \neq i}^{n} \overline{f_j}(r_{i,ref} - r_{j,ref})$$

where n is the number of groups, and $\overline{f_j}$ is the average absolute frequency of the group in the period of time considered.

This average relative growth rate $\overline{r_i}$, can be identified as the selection rate coefficient of the group in the population considered(*48*). The selection rate coefficient is a direct measure of the fitness advantage of emerging variants and is one of the best indicators as to whether a strain will increase in frequency during an outbreak(*49*, *50*).

We can further multiply the selection rate by the mean generation time *(35 days)* to obtain the selection coefficient per generation, which is the relative fitness advantage per transmission generation.

We also estimated the growth rate per serotype to capture whether the individual dynamics were concordant with, or deviated from, what was expected according to their respective group (NVT, PCV7, or PCV13)(Figure 4A-C).

To investigate whether the serotypes fitness changed across upon implementation of PCV7 and PCV13, we tested a range of models. We considered a model without any shift of fitness, a model with a single shift in 2009 and a model with two growth rates represented by a shift in fitness in both 2009 and 2011. Model comparison was done using the Watanabe-Akaike information criterion (WAIC) implemented in the loo package(*51*). Further, we also tested for a potential delay between implementation of PCV7 and the change in fitness. Model comparison is presented in Figure S22.
The model performed best when the fitness switch was in 2009 (the initial year of vaccine implementation). We used this model in the main text.

To fit the model, we used Hamiltonian Monte Carlo as implemented in R-Stan(*52*). We used a Poisson likelihood to fit the observed proportion of sequences that were of each category and the total number of isolates in that year as an offset. The model estimated the proportion of isolates that were of each type at the start of the dataset and the fitness parameters.

#### *Antimicrobial Resistance*

We then used the model to capture the decreasing proportion of penicillin resistance in the population. To do this we incorporated the dynamics of vaccine type (VT; PCV7 and PCV13) and NVTs in the population, and the respective proportion of penicillin resistance within them.
We used the same approach described above to model the proportion of VTs, $f_{vt}$, and NVTs, $f_{nvt}$. We then model the proportion of strains which were and were not penicillin resistant within each group, either VT, $f_{amr|vt}$, or NVT, $f_{\mathrm{amr|nvt}}$. We estimate the fitness of resistance in each.
We then fit the model to all four groups, $f_{vx,amr}$ (Resistant NVT, Susceptible NVT, Resistant VT, and Susceptible VT).
Equation 13.

$$f_{vx,amr}(t) = f_{vx}(t) \cdot f_{amr|vx}(t)$$

Where $f_{vx}$ is the proportion of either VT or NVT in the population and $f_{amr|vx}$ is the proportion of penicillin resistance within each of those groups. We then derive the proportion of penicillin resistance overall in the population by summing the proportion of VT resistant, $f_{vt,amr}$, and NVT resistant, $f_{nvt,amr}$, strains.

### Estimating Impact of Fitness on Migration

We included the calculated average relative growth rates in the simulated branching epidemic to calculate the mean number of municipalities visited, distance travelled, and probability of being in the home municipality over 5 years. We report uncertainty at one standard deviation from the mean. We repeated this incorporating the post-PCV selection coefficients for NVTs and VTs and estimated the relative increase in the number of municipalities visited for NVT serotypes compared to PCV serotypes after 2 years of transmission.

### Fitness Model Carrying Capacity Sensitivity

Our fitness model assumes constant fitness over time. This specifically means that if a serotype is found to be fitter than the rest of the population, we expect it to replace the whole population after some time. However, the currently best supported model proposes that negative frequency dependent selection drives pneumococcus population dynamics (*53*), whereby lower frequency genes become more fit.

To test the impact of our assumption on our fitness estimates, we performed a sensitivity analysis. We introduction minimum and maximum carrying capacities in our logistic model. In equation 14, we let the relative frequency $f_{i,ref}$ go to a maximum of $K_{max}$, and in equation 15, a minimum of $K_{min}$:

Equation 14

$$f_{i,ref}(t) = \frac{K_{max}}{1+\left(\frac{K_{max}-f_{i,ref,0}}{f_{i,ref,0}}\right)\exp(-r_{i,ref}\cdot t)}, \text{ if } r_{i,ref} > 0$$

Equation 15

$$f_{i,ref}(t) = 1 - \frac{1-K_{min}}{1+\left(\frac{(1-K_{\min})-(1-f_{i,ref,0})}{(1-f_{i,ref,0})}\right)\exp(r_{i,ref}\cdot t)}, \text{ if } r_{i,ref} < 0$$

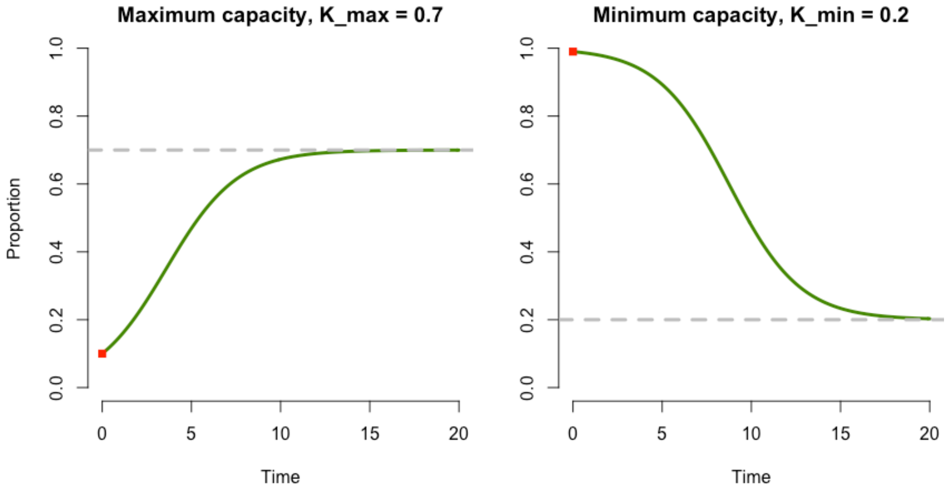

with $K_{min} \leq K_{max}$, and $K_{min} \in [0,1], K_{max} \in [0,1]$.

We considered a range of values for $K_{min}$ (0.1, 0.05 and 0) and $K_{max}$ (0.9, 0.95 and 1). We tested the impact of this carrying capacity has on both the vaccine status model (Figure S23), and antimicrobial resistance and vaccine status model (Figure S24). In each case, we compare the fitness estimates obtained. We found the fitness estimates to be robust to variable carrying capacities across those tested implying our assumption does not impact this framework.

### Code Availability

All scripts for analysis are available on GitHub at:

https://github.com/sophbel/Analysis_GeographicMobility_pneumoPaper

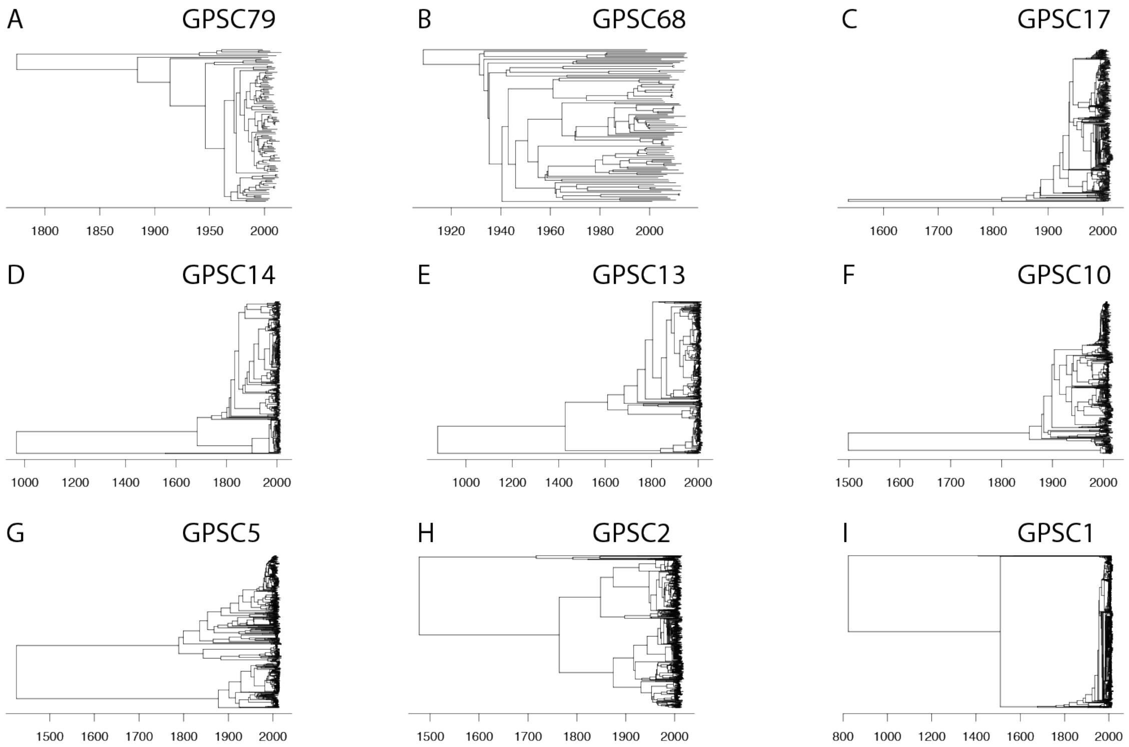

**Fig. S1A-I. Time Resolved Trees for Dominant GPSCs.** Trees are recombination masked, aligned to a reference for each GPSC. Time resolution was performed using BactDating.

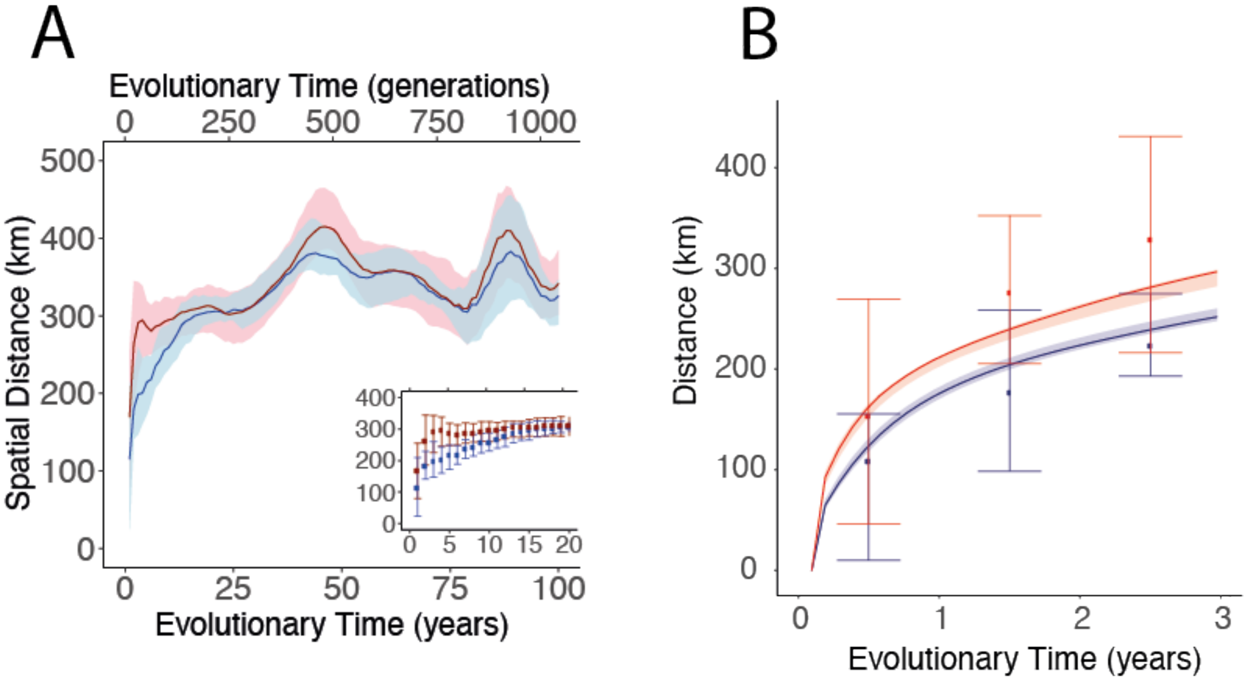

**Fig. S2.** A) Mean geographic distance (km) across rolling 20-year divergence time windows between pairs for disease isolates alone (red) and for all isolates (blue). Subset to 20 years only as an inset. B) Mobility model fit (lines) against data (points) for disease alone (red) and all isolates (blue). This is a subset from Figure 1D.

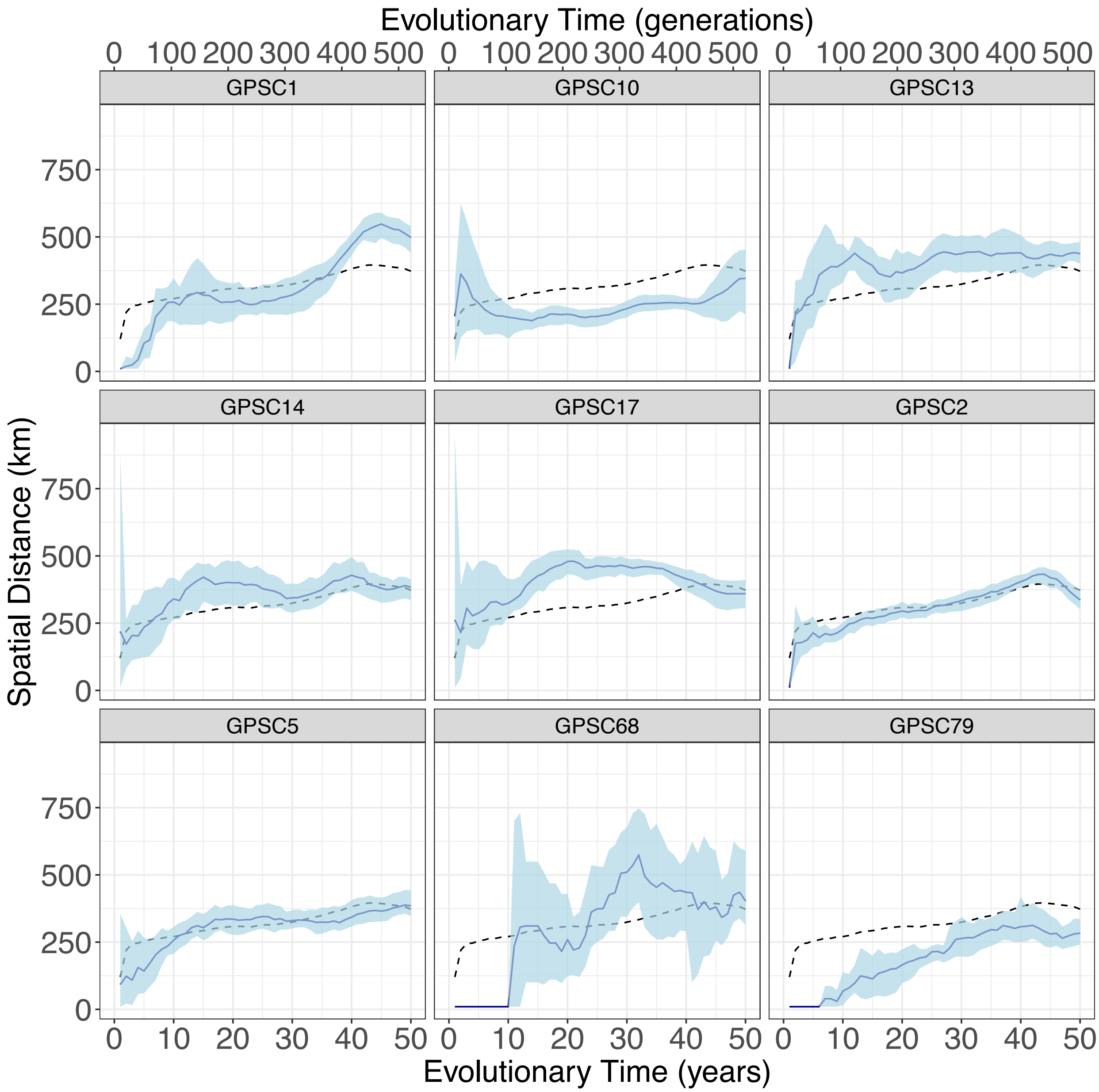

**Fig S3**. Mean geographic distance (km) across rolling 20-year divergence time windows between pairs for each of the 9 dominant GPSCs (light blue) against the overall line from the aggregated data (purple dashed).

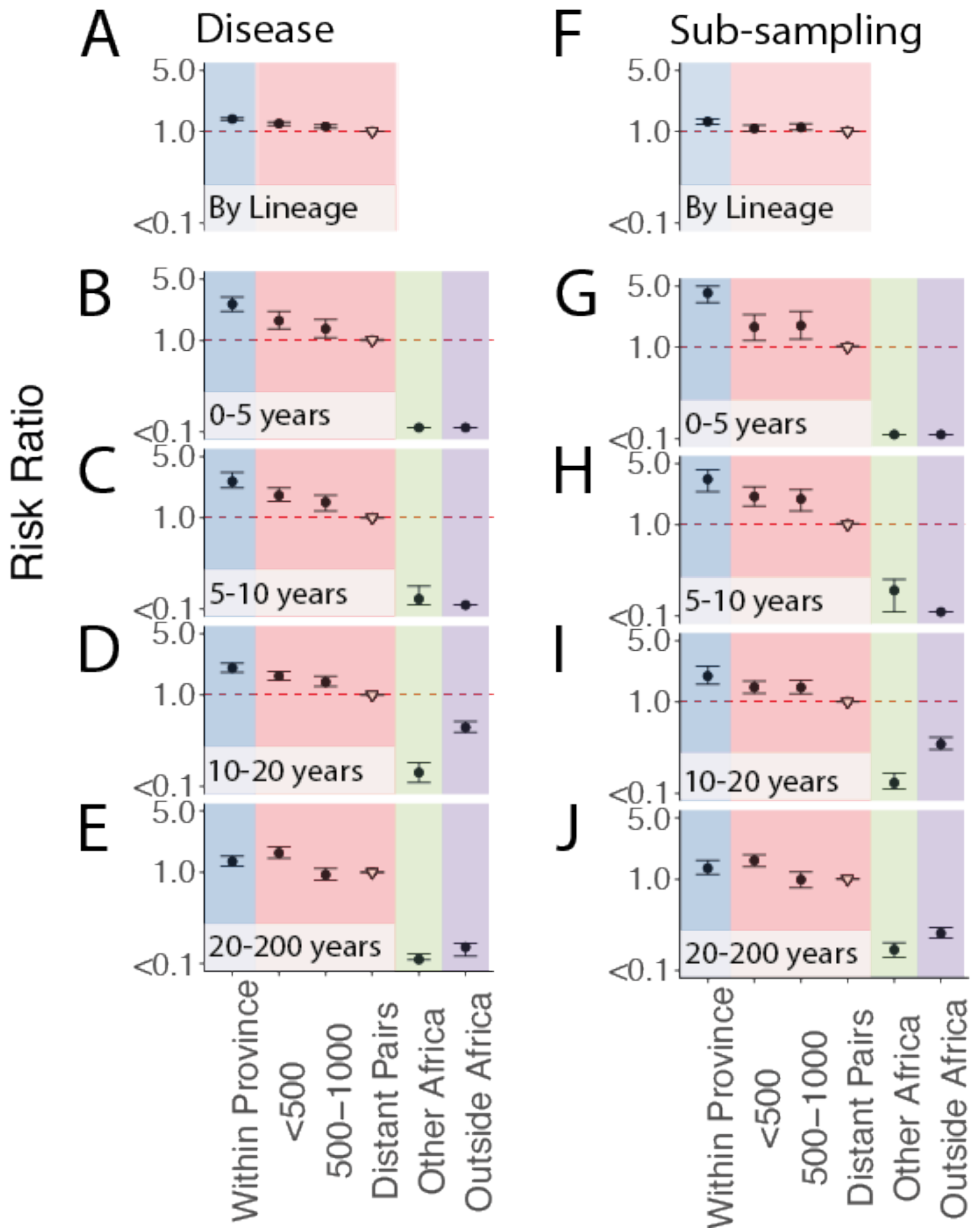

**Fig. S4 Risk ratio framework to determine geographic structure for only disease isolates. (A-E)** A) Risk ratio of being the same GPSC within a province (blue), between different provinces over increasing distance (red), compared to geographically distant pairs (>1000km) (reference). (South Africa; N=6910) B) Risk Ratio of having tMRCA 0-5, C) 5-10, D) 10-20, or E) 20-200 years ago within South African provinces (blue), across larger distances within South Africa (red), from South Africa to other countries in Africa (green), and from South Africa to countries outside of Africa (purple). All plots use a reference of pairs which are from distant provinces in South Africa (open triangle Risk ratio framework to determine geographic structure for isolates sub-sampled to 300. (F-J) are the same as above.

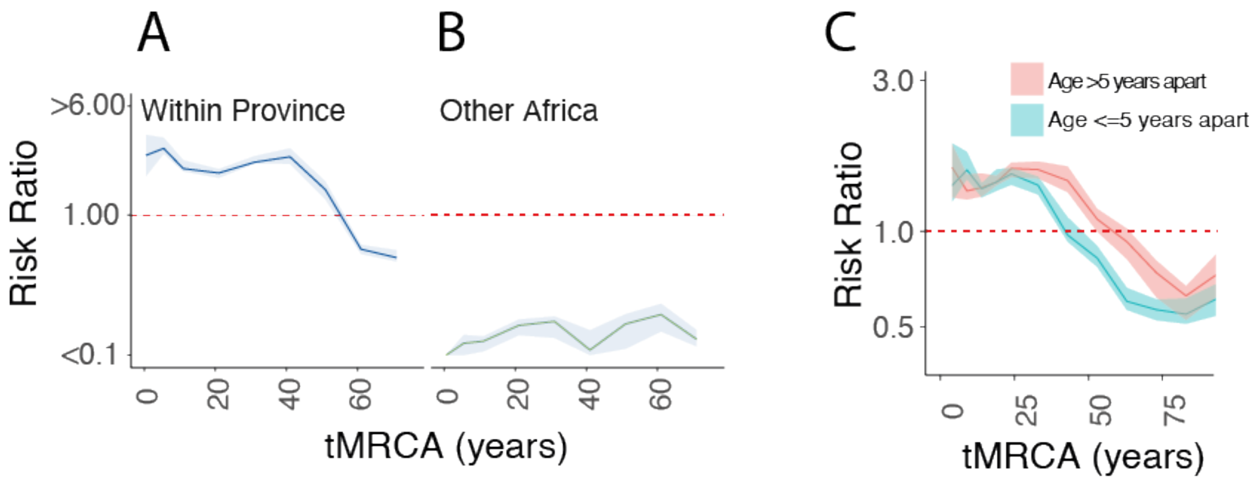

**Fig. S5.** Relative Risk of similarity over rolling 20-year windows of divergence times for pairs isolated A) within the same South African province compared to pairs from distant provinces in South Africa (>1000km apart), B) and for pairs where one isolate is from South Africa and the other is from elsewhere in Africa. C) Relative Risk of similarity over 50-year rolling window of divergence time for pairs isolated from people whose age at collection time was <=5 years apart (blue) or >5 years apart (pink).

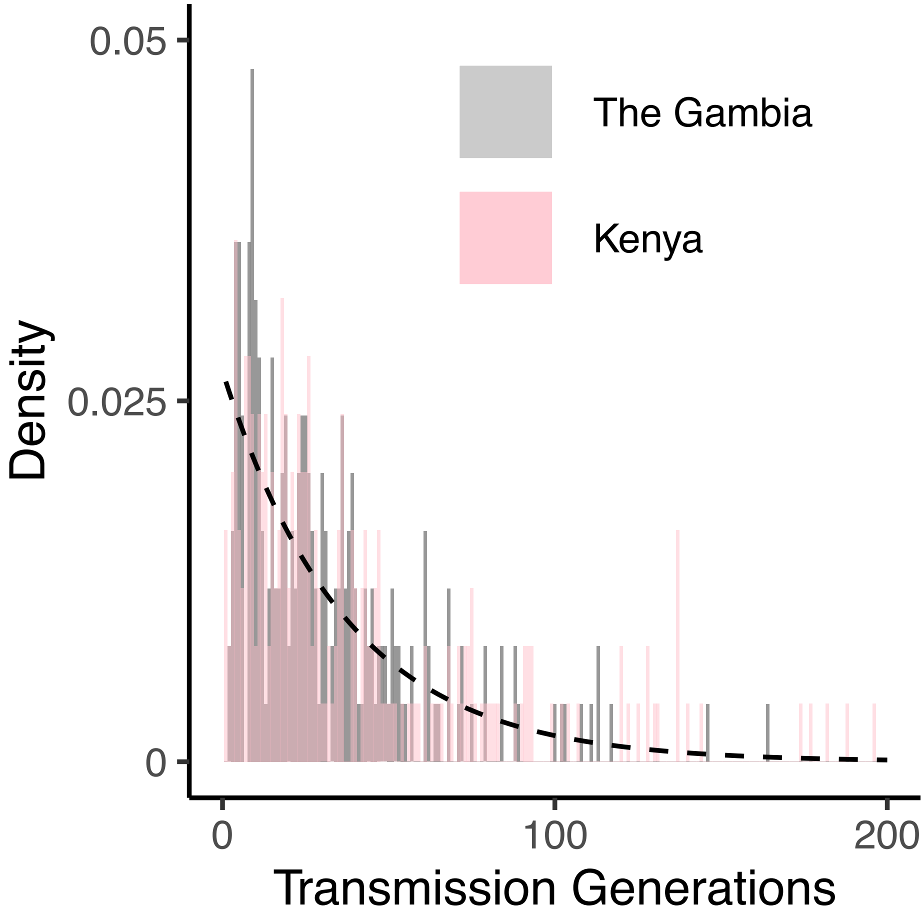

**Fig. S6.** Histogram of simulated transmission generations from The Gambia (grey), and Kenya (pink), overlayed with a gamma distribution curve scaled to the mean of these with a shape of 1 and scale of 0.096 (dashed black).

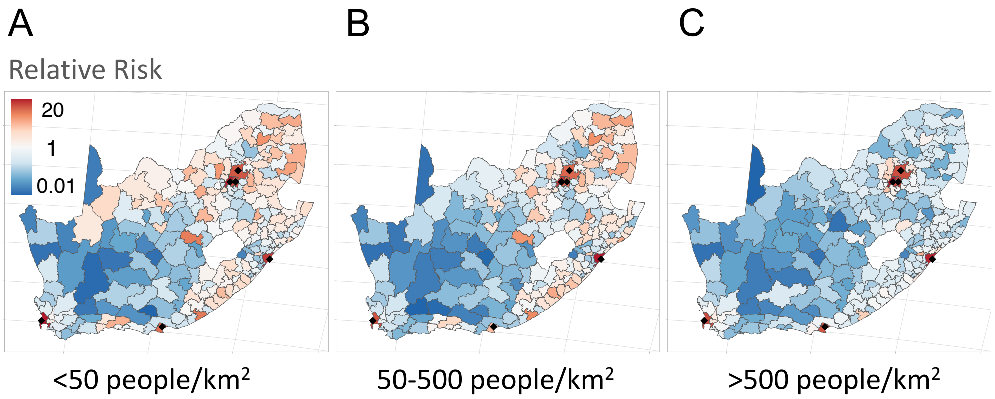

**Fig. S7. Relative risk of a pneumococcal strain being in each of municipality after 1 year of transmission.** Sequential transmission chains starting in municipalities with A) <50 people/km2, B) 50-500 people/km2, or C) >500people/km2

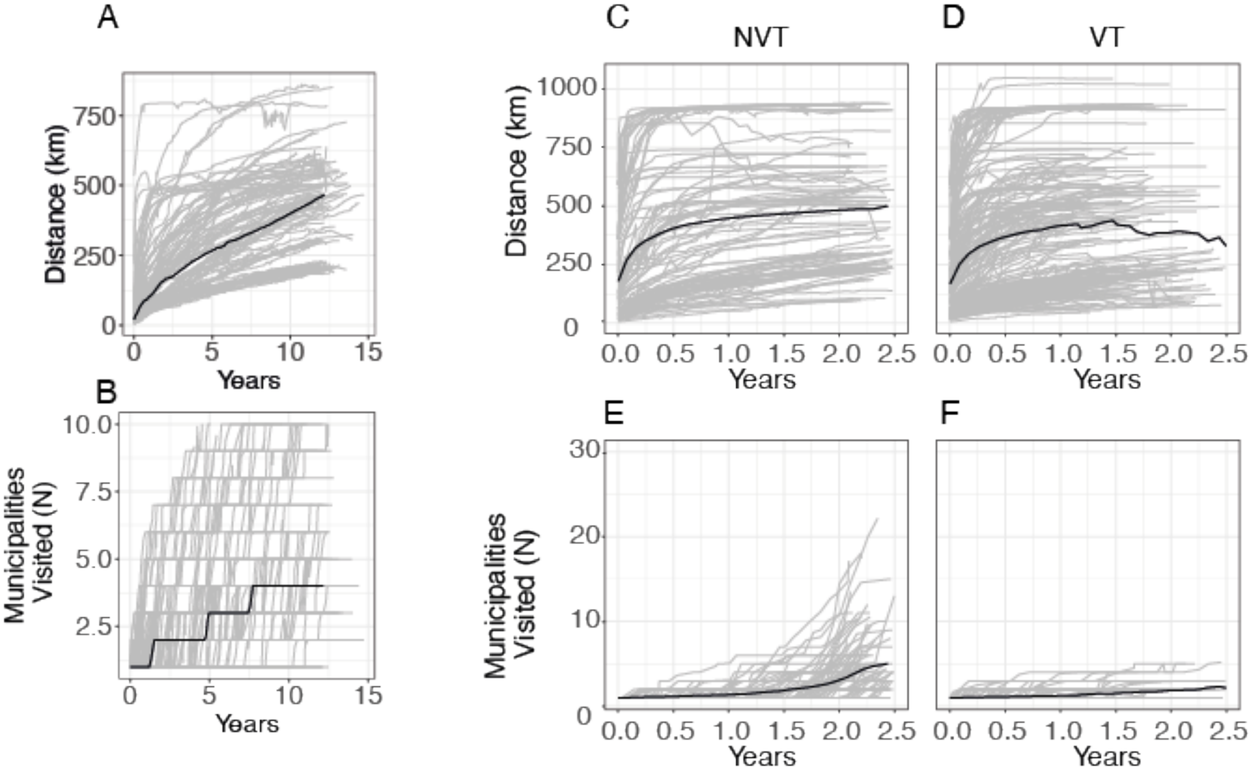

**Fig. S8.** Branching epidemic simulations adjusting the effective reproductive number ($R_{eff}$)A & B) using a $R_{eff}$ of 1 and a Poisson distribution and C-F) adjusting the $R_{eff}$ by the relative fitness estimated using the fitness model in the post-PCV era. C & E) for NVT serotypes, and D &F) for VT serotypes utilizing the estimated fitness for the metrics A,C,D) distance traveled per simulation, B,E,F) number of unique municipalities visited per simulation.

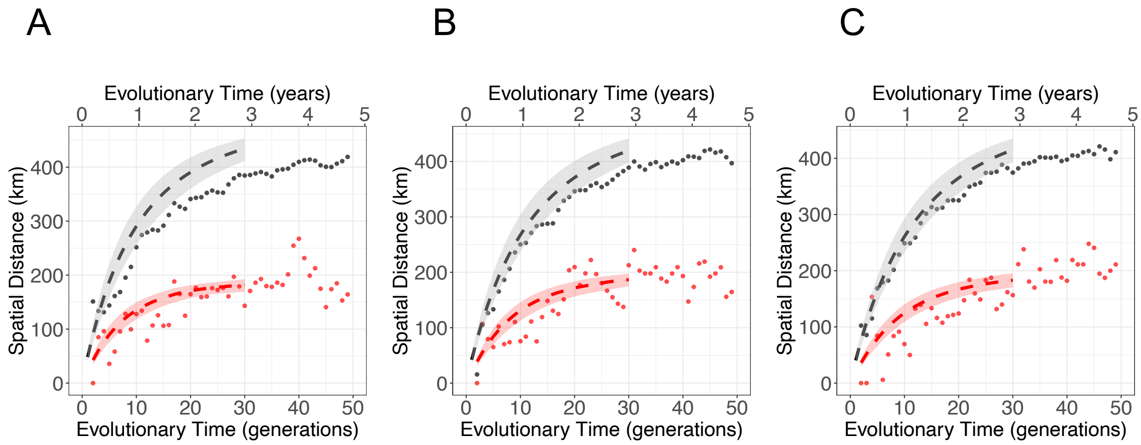

**Fig. S9. Independent replicated simulations (A-C) for model performance testing.** Simulated epidemic data (black dots), down-sampled simulations (red dots), model fit to down-sampled data (red line), removing the sampling probability the model recapturing the true epidemic (black dot).

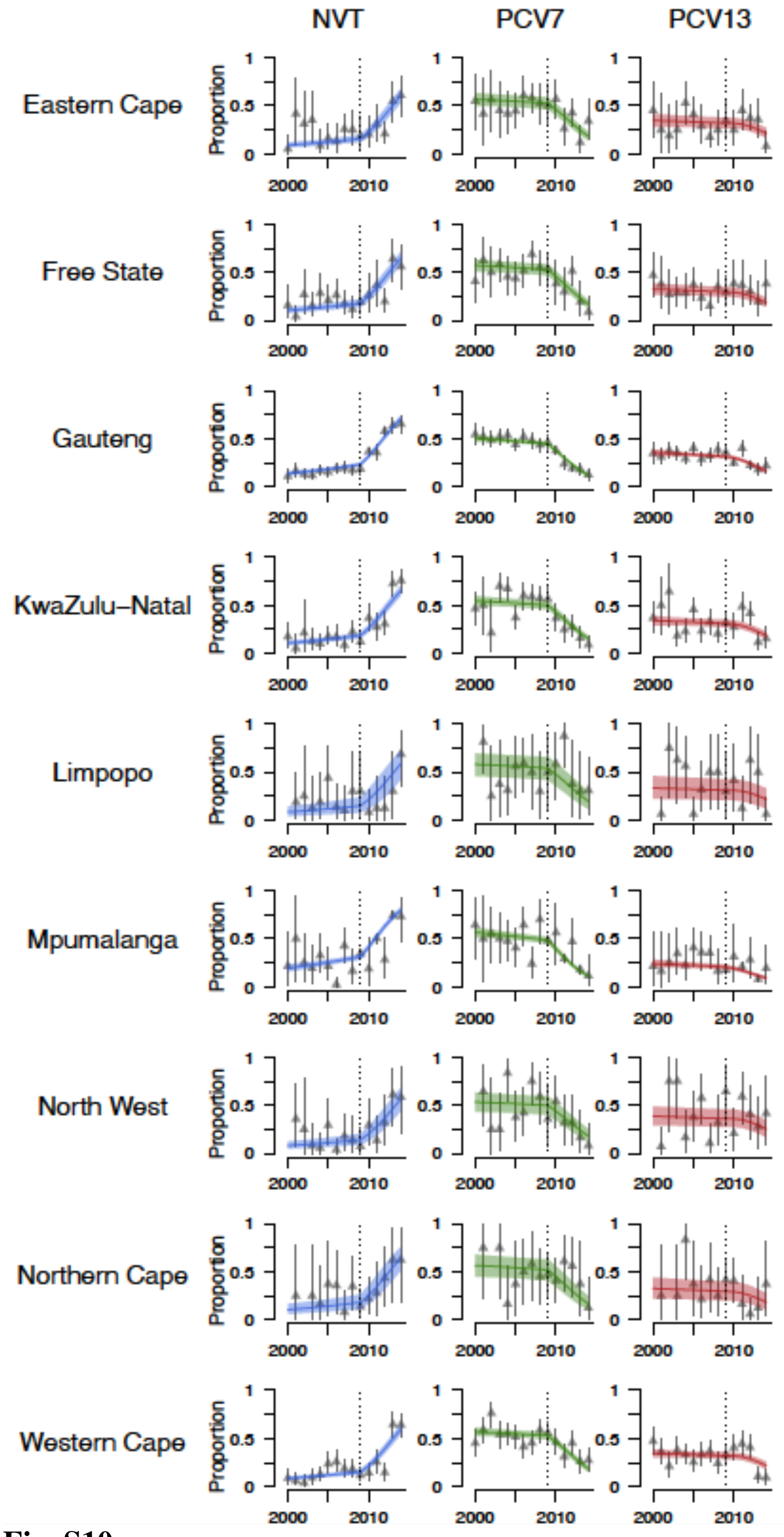

**Fig. S10.**
The proportion of groups across provinces over time from (2000-2014) for NVT serotypes (blue), PCV7 serotypes (green), and PCV13 serotypes (excluding those in PCV7) (red) from the data. The model fits overlay the data including a fitness change in 2009 as indicated by the vertical dashed line.

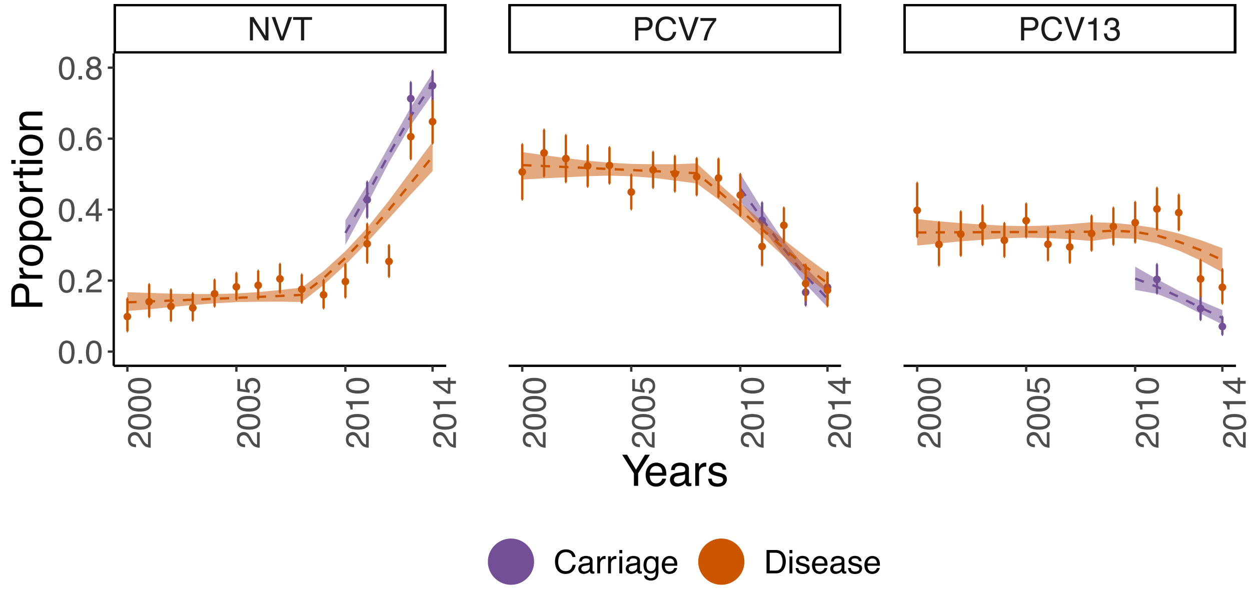

**Fig. S11** Proportions of four groups in the population (points) when looking at only isolates collected from carriage (purple) and isolates collected from disease (orange) overlayed by model fit (lines).

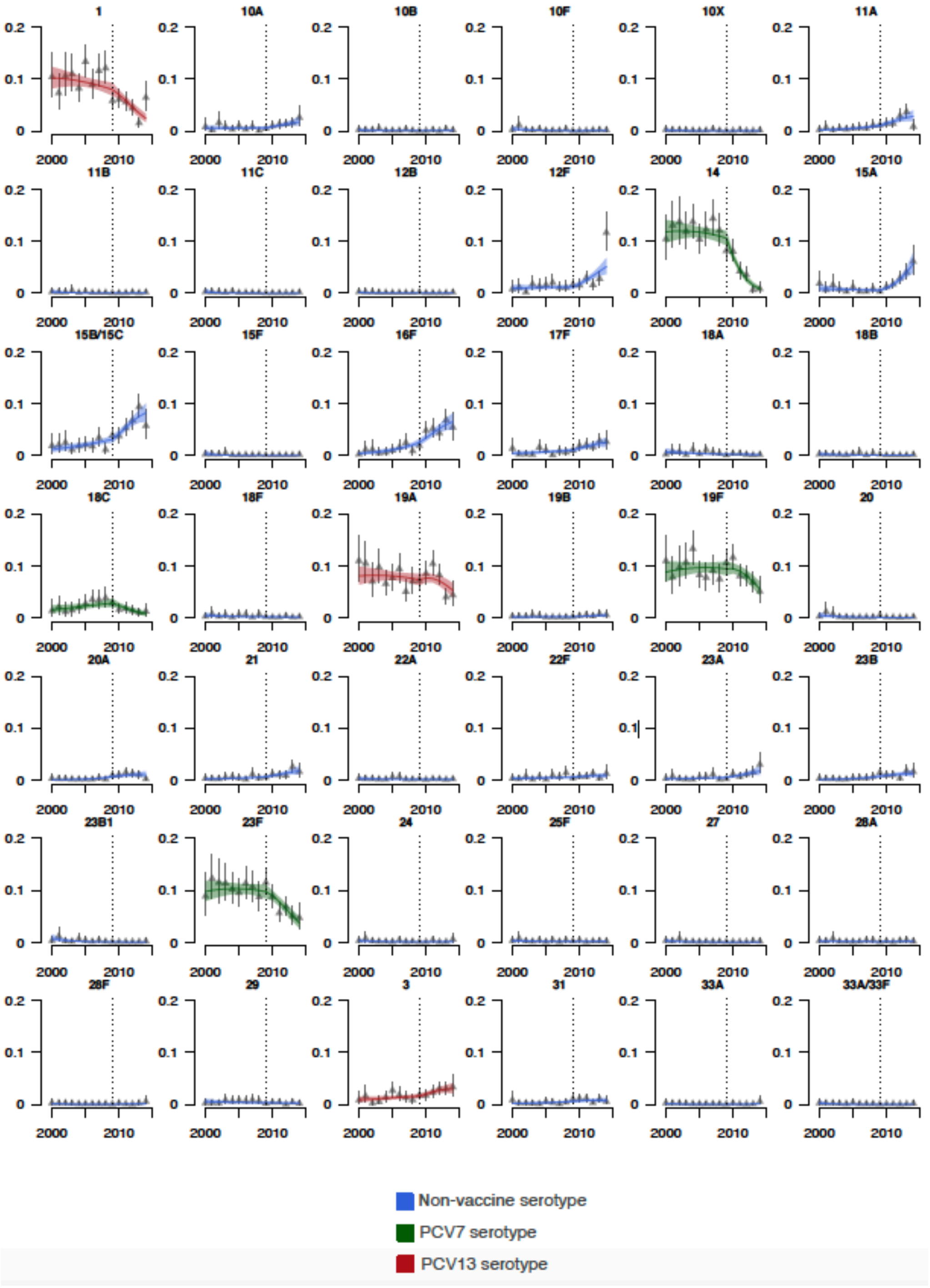
1
10A
10B
10F
10X
11A
11B
11C
12B
12F
14
15A
15B/15C
15F
16F
17F
18A
18B
18C
18F
19A
19B
19F
20
20A
21
22A
22F
23A
23B
23B1
23F
24
25F
27
28A
28F
29
3
31
33A
33A/33F
0.2
0.1
0
2000
2010
Non-vaccine serotype
PCV7 serotype
PCV13 serotype

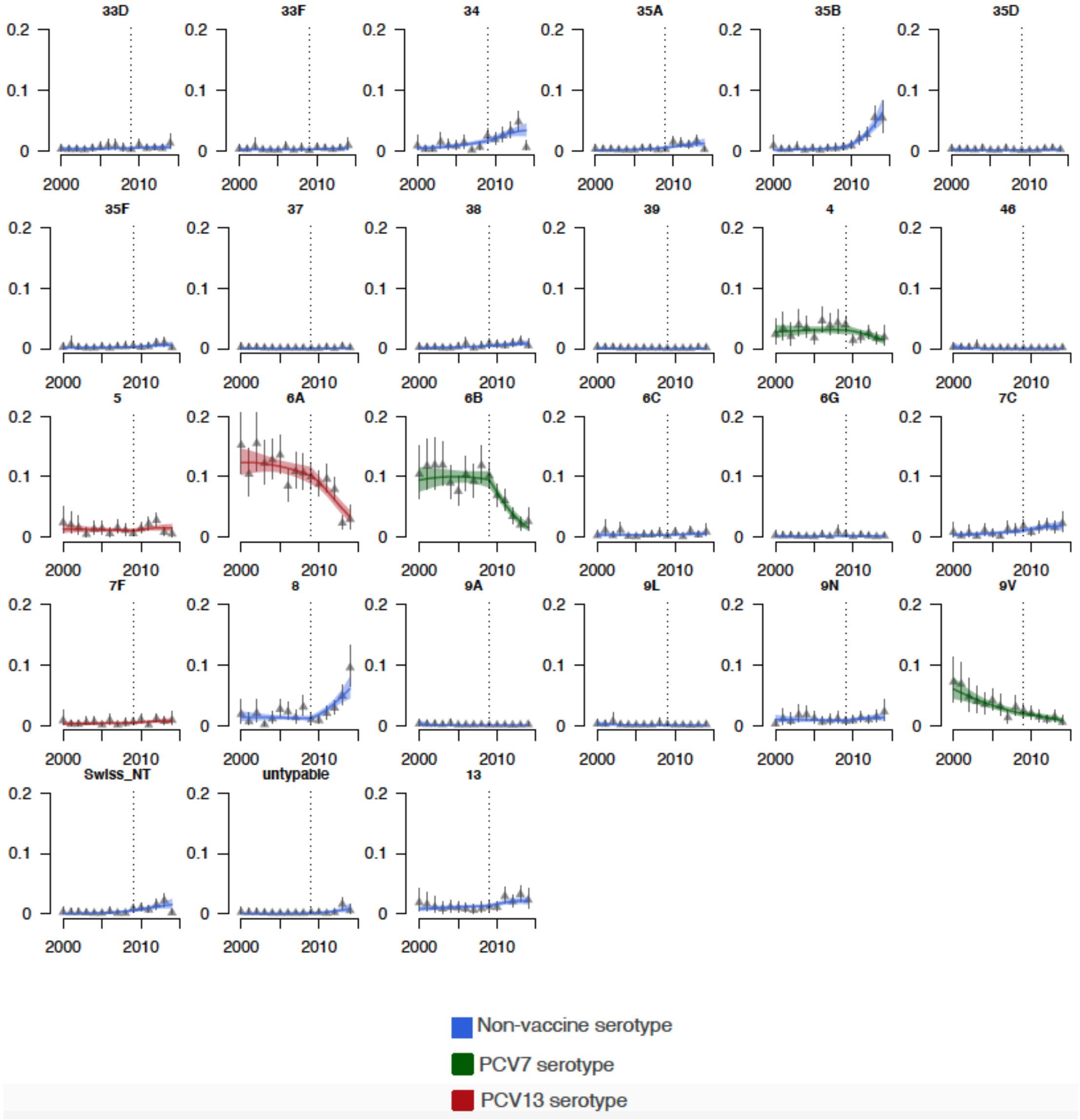

**Fig. S12.** Fits by serotypes colored by whether they are included in PCV7 (green), additionally included in PCV13 (maroon), or not included in any vaccine (blue). Estimating the growth rate prior to PCV7 implementation (2009), and after implementation indicated by the vertical dashed line.

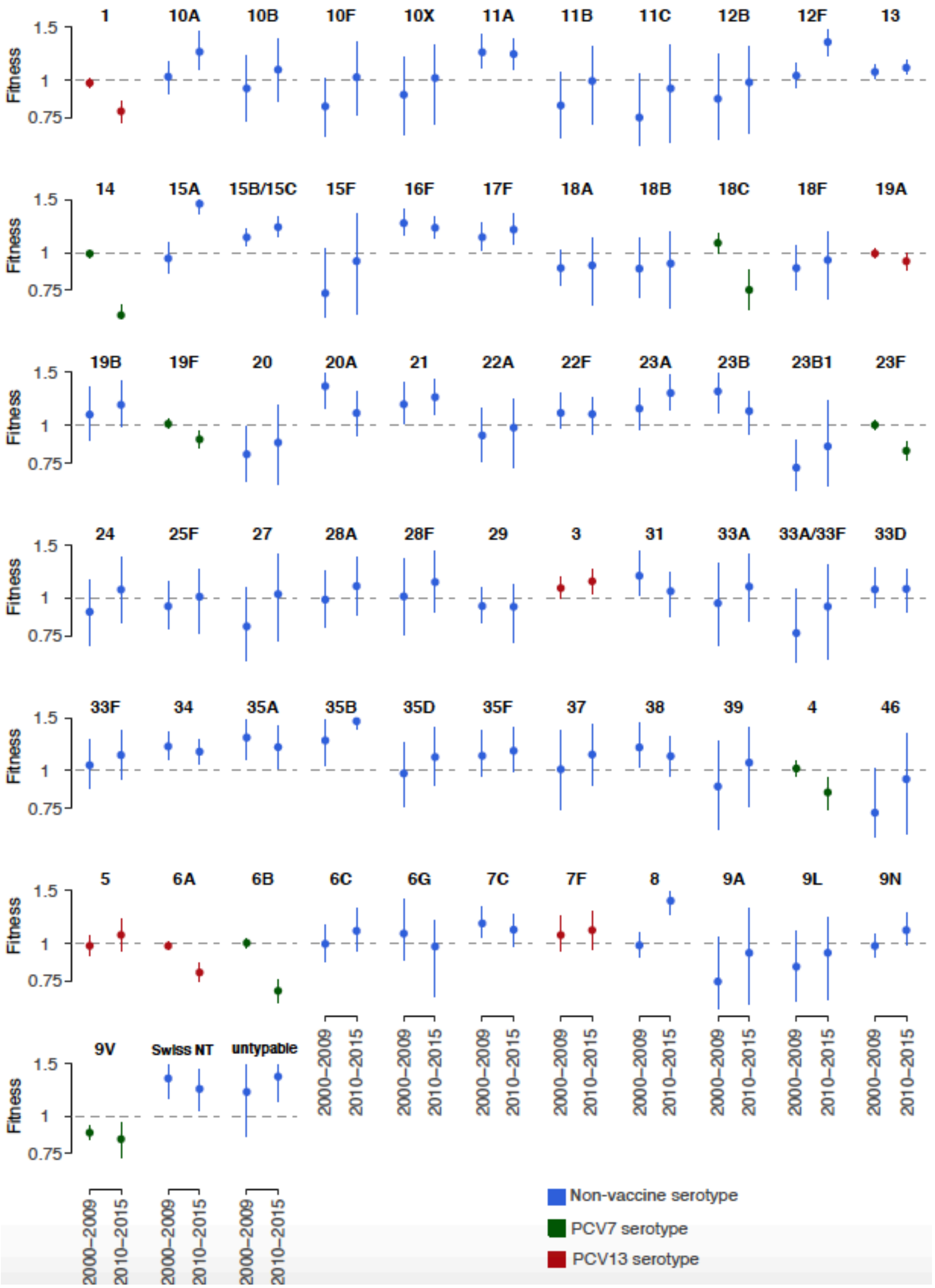

Fig. S13. Fitness estimate for each serotype. Estimates for years prior to PCV7 implementation (2009), and after PCV7 implementation. Colored by whether they are included in PCV7 (green), additional types included in PCV13 (maroon), or not included in any vaccine (blue).

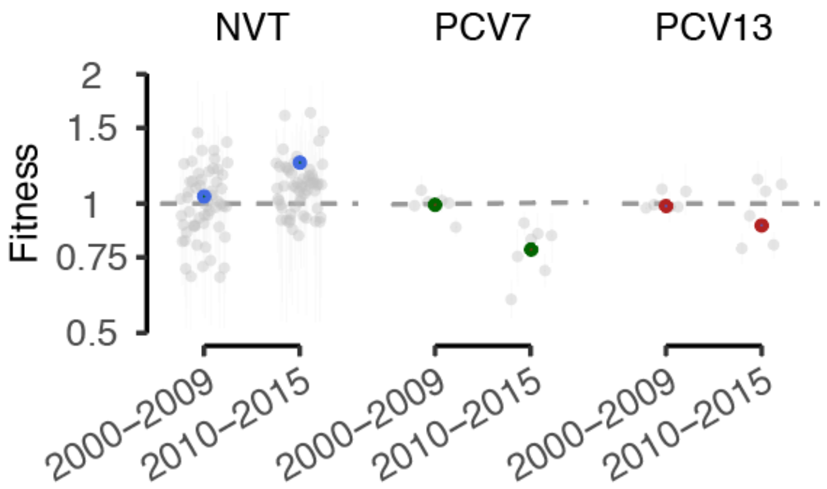

**Fig. S14.**
Fitness estimate for each serotype (grey), superimposed by group.

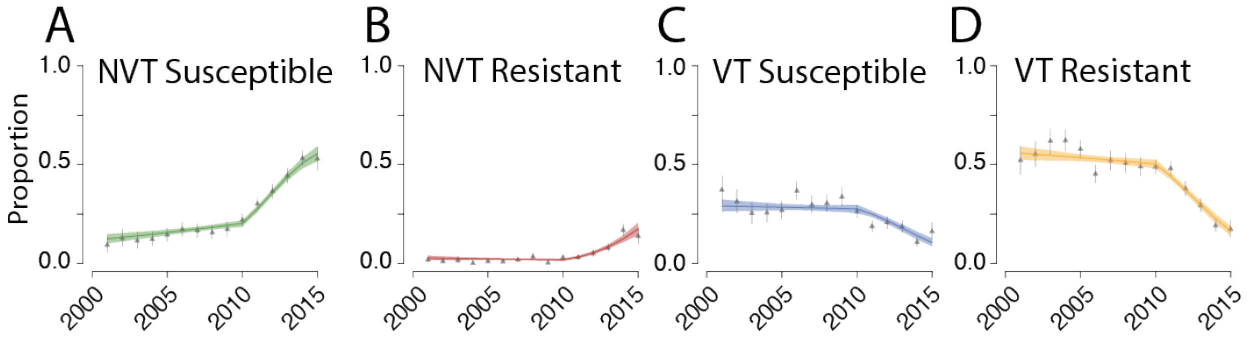

**Fig. S15** Data fits for model accounting for proportions and fitness over time in four groups, A) NVT-penicillin susceptible (green), B) NVT-penicillin resistant (red), C) VT-penicillin susceptible (blue), and D) VT-penicillin resistant (yellow). Estimating a shift in fitness in 2009.

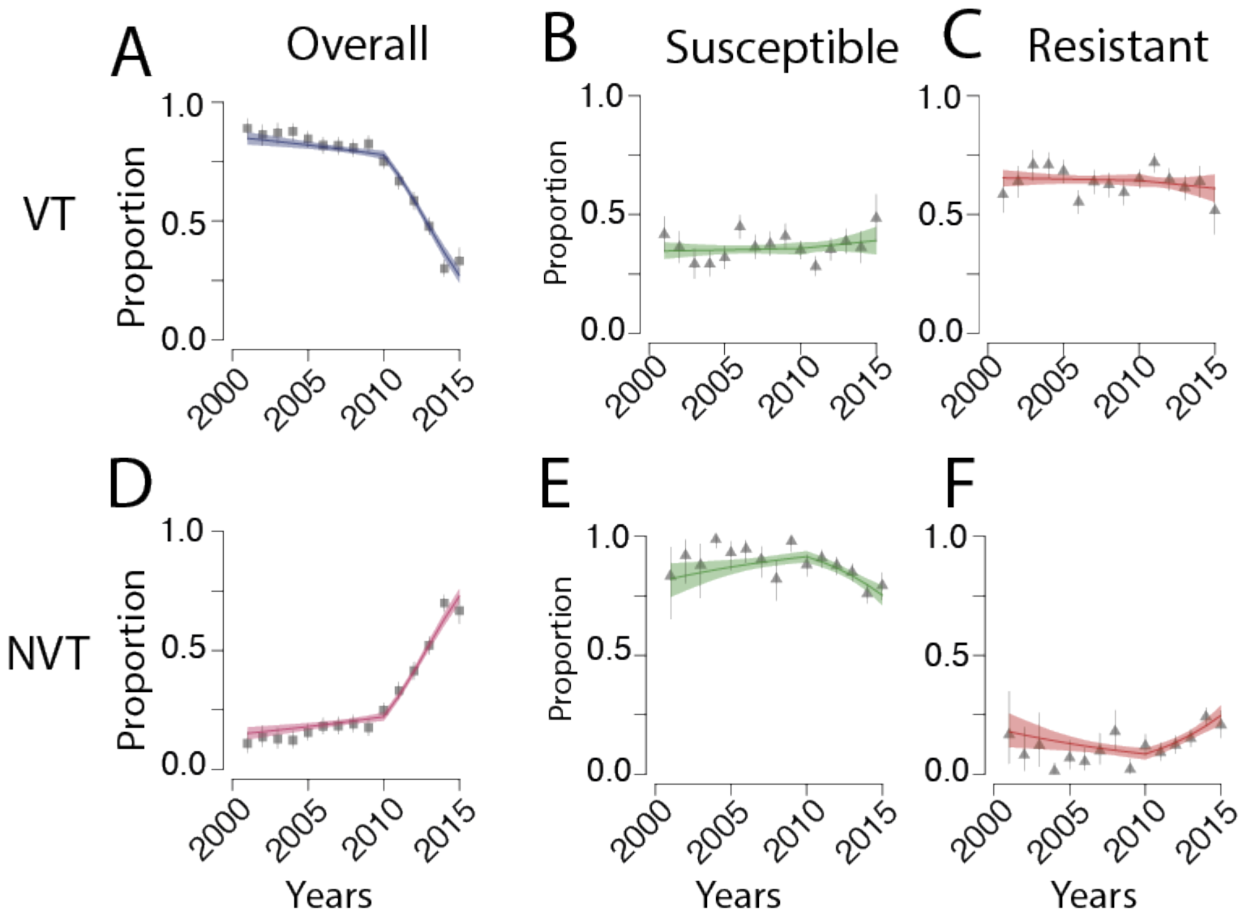

**Fig. S16.** Fits for the individual proportions over time in the vaccine status & AMR model. Proportion of each group changing over time for VTs (A-C), and NVTs (D-F). A) The proportion change of VT over time included with model fits and D) is the proportion change of NVT over time. Additionally, we include the proportion of (B&E) penicillin susceptible (green) and (C&F) penicillin resistant (red) isolates in each of those groups (NVT and VT) over time with model fits.

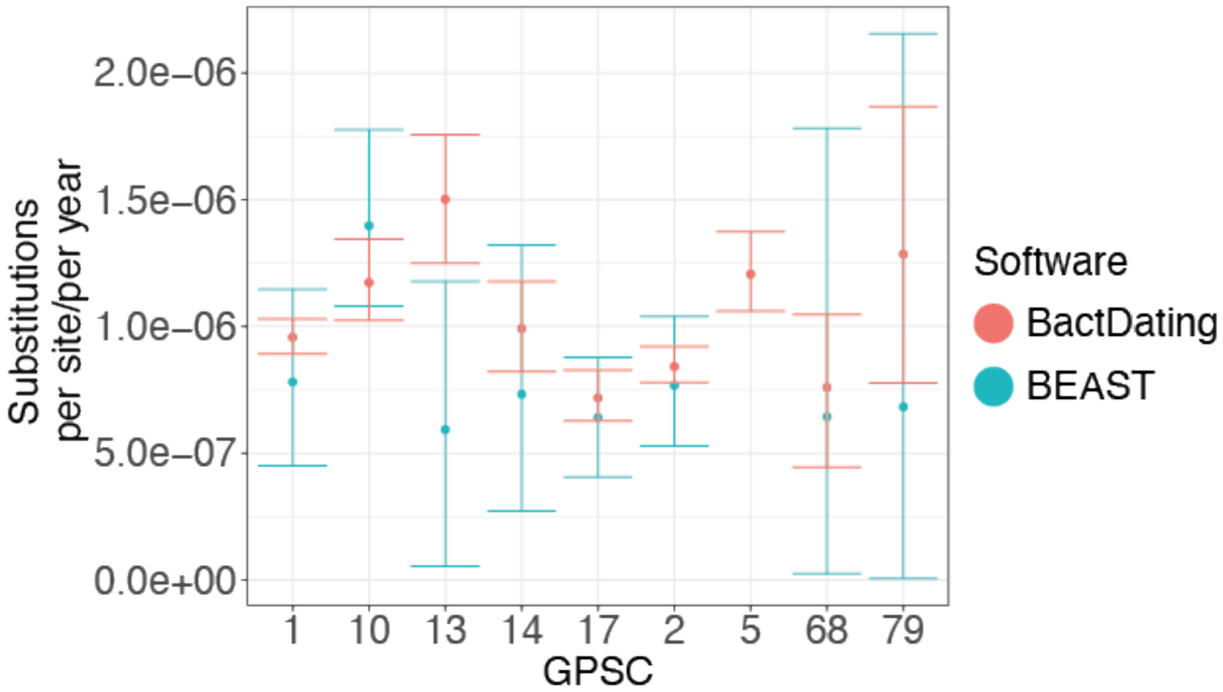

**Fig. S17. BEAST/BactDating comparison.** Comparing the substitutions per site per year from a BactDating relaxed clock model with recombination masked by Gubbins and BEAST relaxed clock model for each of the 9 GPSCs. BEAST was intractable for the diversity in GPSC5.

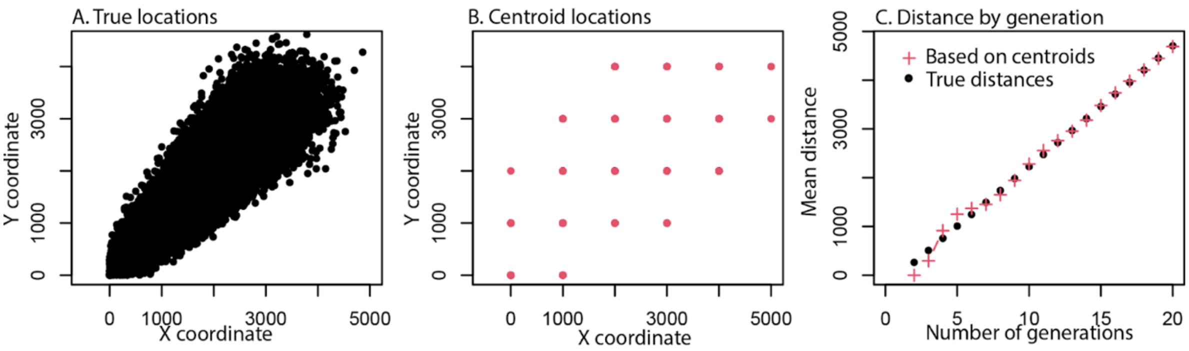

**Figure S18. Comparing centroid distance estimates to true distances.**

To demonstrate the suitability of using centroid distances, we simulated a spatial transmission process for 1000 separate chains where at each generation a daughter point is placed at a randomly located location 350m in each of the x and y direction. A) This is repeated over 20 generations. B) We then identify the 'centroid' of each case based on the closest coordinate rounded to the nearest kilometre C). We then calculate the total distance covered for both the true distance (black) and the centroid distances (red).

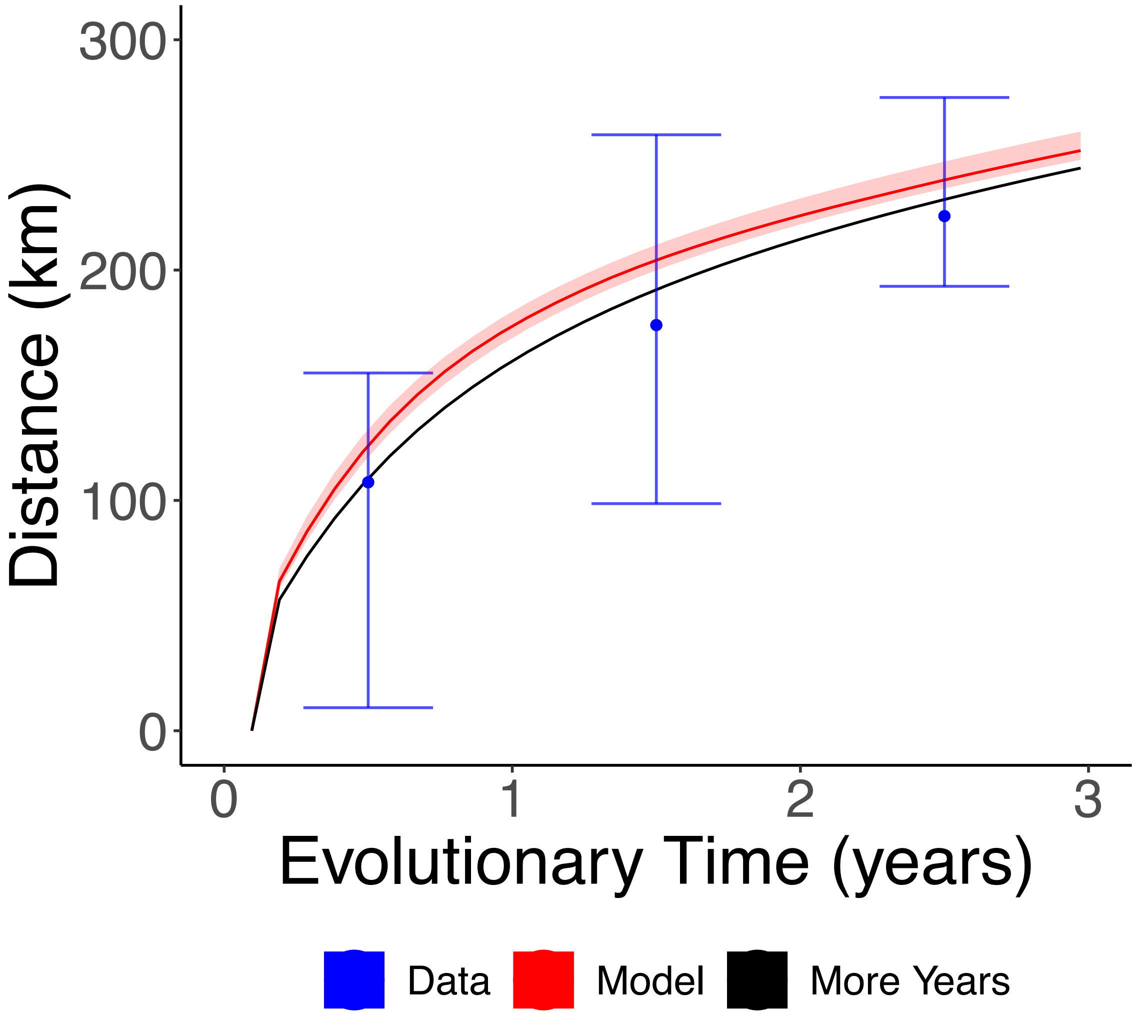

**Fig. S19. Fitting the mobility model to 15 years of evolutionary distance.** Fitting the probabilistic mobility model to pairs of genomes which are 15 years divergent from their MRCA (black), compared against the 10 years we used (red), and the data (blue).

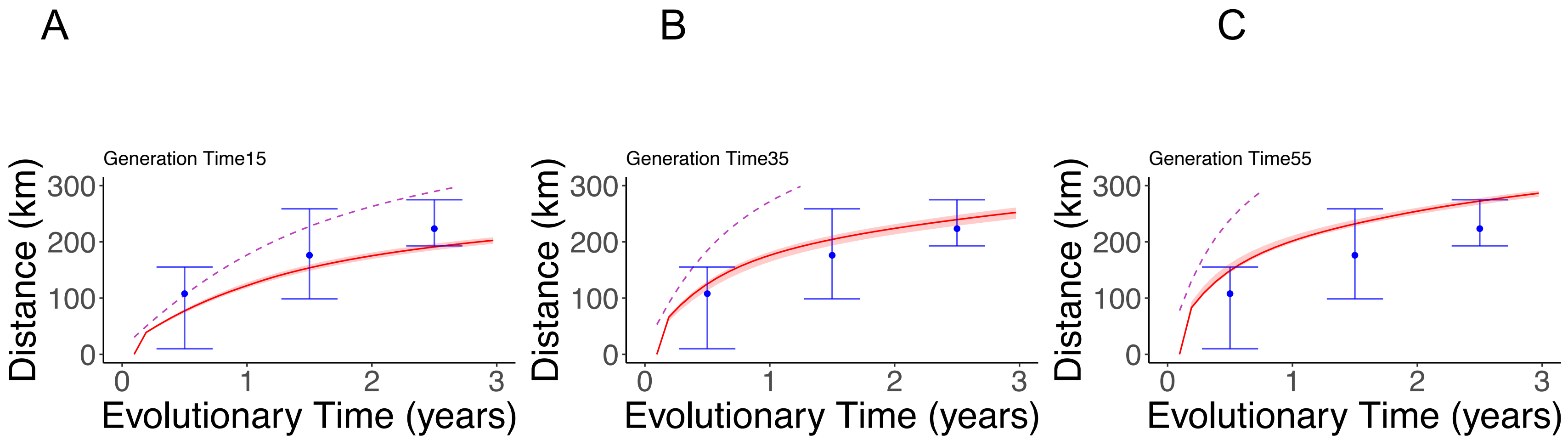

**Fig. S20. Comparing model fit with variable generation time estimates.**

Assessing the mean geographic distance per evolutionary time in years for the data (blue), including the sampling probability (red), excluding the adjustment for sampling probability (purple dashed) for A) generation time of 15 years, B) generation time of 35 years, and C) generation time of 55 years.

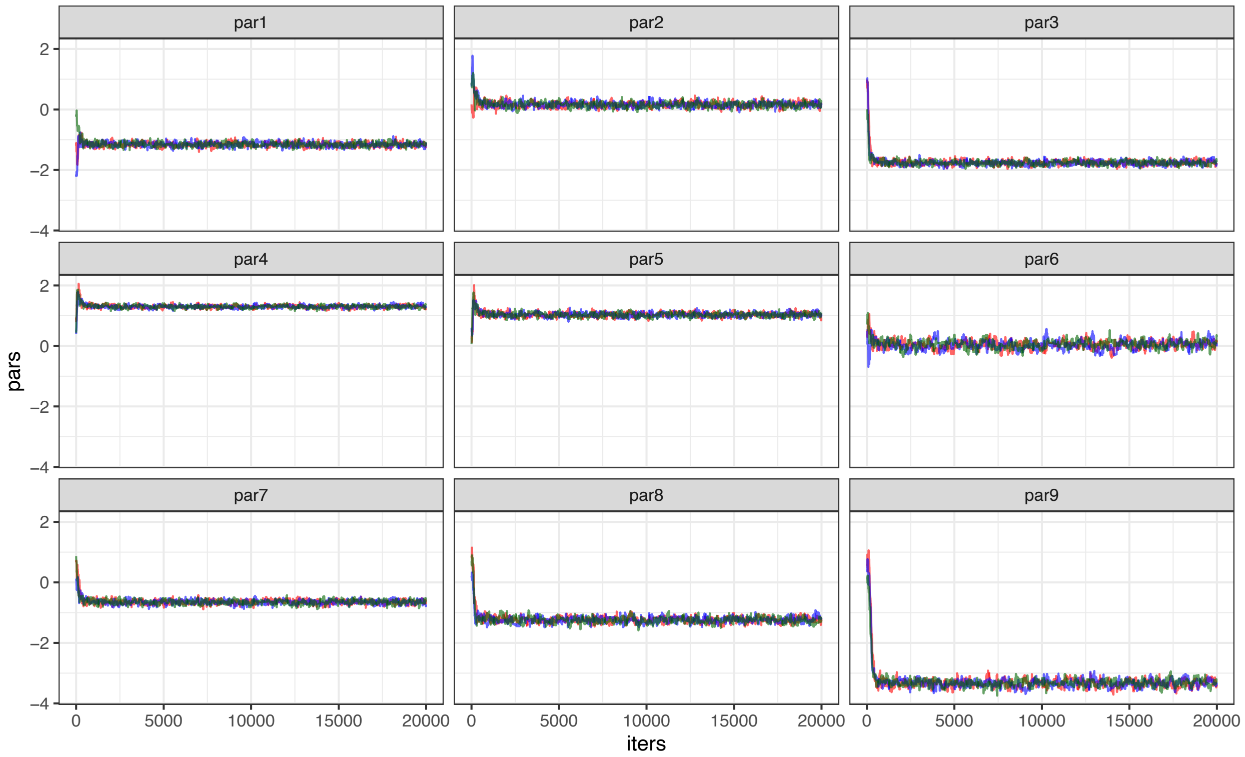

Figure S21. Converging chains across 20000 simulations for 9 parameters estimated in the model.

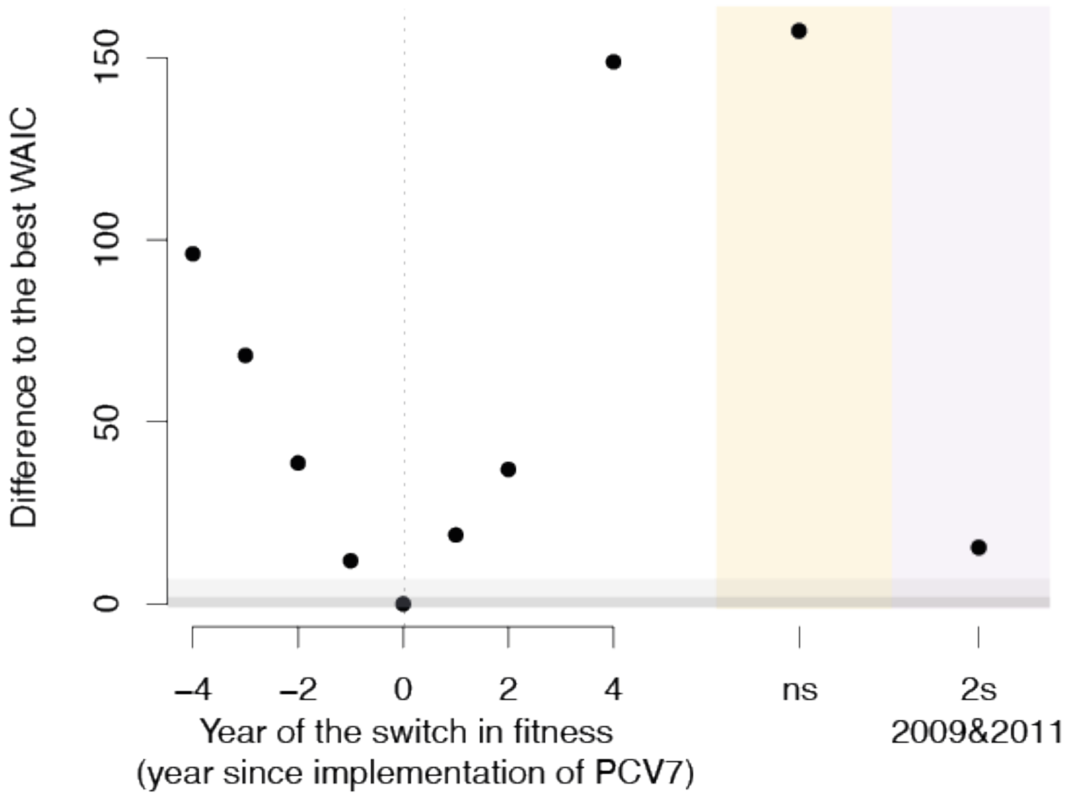

**Fig. S22. Testing year of fitness switch for the logistic growth -fitness model.**

Adjusting the year of the fitness switch in the model fitting to vaccine status. The difference to the best WAIC (2009) is on the y-axis where the year of fitness switch relative to 2009 is on the x-axis. Further we test no fitness switch (ns; yellow) and testing the impact of including two fitness switches in 2009 (PCV7 implementation) and 2011 (PCV13 implementation) (purple).

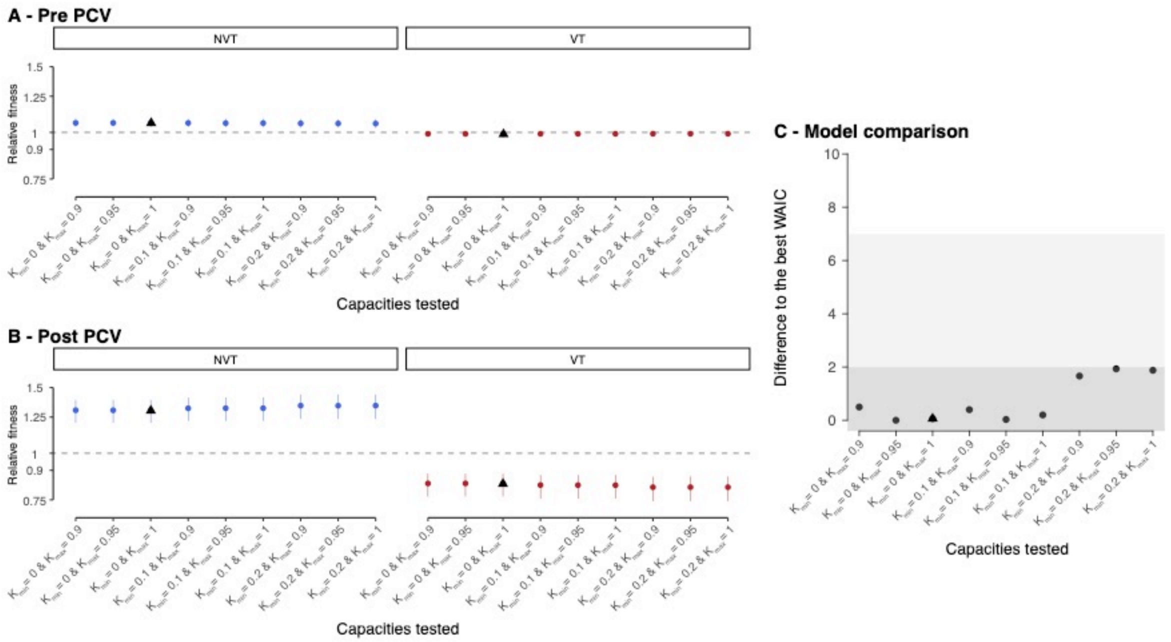

Figure S23. **Fitness estimates with varying carrying capacity across VT and NVT serotypes** A) pre-PCV, B) post-PCV introduction, C) and testing the difference to the best WAIC for each. The black triangle indicates the carrying capacity utilized in the main model.

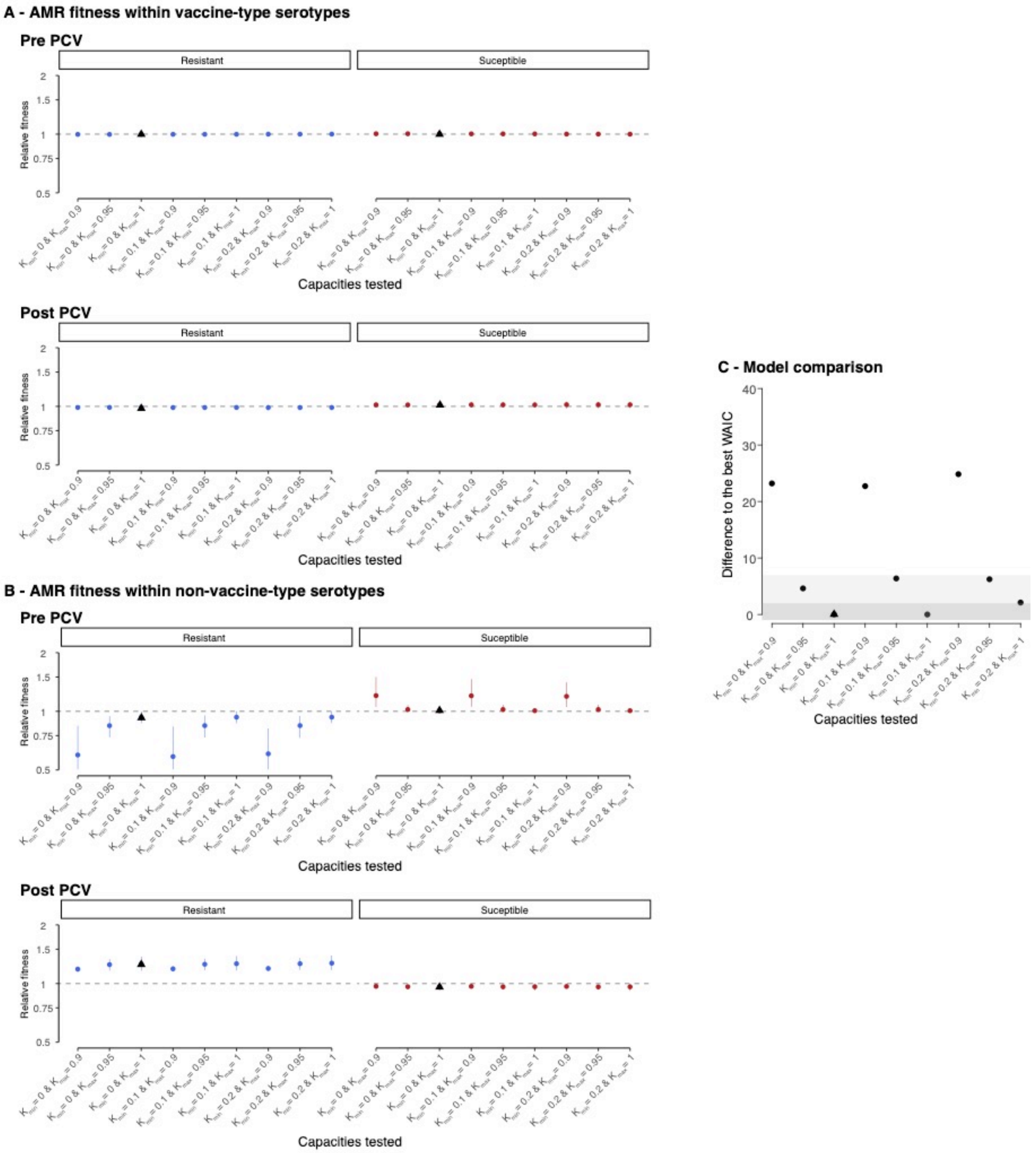

Figure S24. **Fitness estimates with varying carrying capacity** for penicillin resistant isolates across A) VT serotypes and B) NVT serotypes where the top row for each is the relative fitness pre-PCV across carrying capacities. and the bottom row is post-PCV introduction relative fitness estimates. C) Testing the difference to the best WAIC for each. The black triangle indicates the carrying capacity utilized in the main model.

**Table S1. Summary table of isolates from the 9 provinces of South Africa including the collection years, total number, pre and post-PCV, isolates from patients with invasive pneumococcal disease (IPD), isolates which are not included in the vaccine (NVTs), and dominant GPSCs.**

| Province | Collection Years | Total | pre-PCV7 | post-PCV | IPD | | GPSCs | NVT Serotypes | | pre-PCV NVT | | post-PCV [illegible] | |
|---|---|---|---|---|---|---|---|---|---|---|---|---|---|
| | | N | | | N | % IPD | N | N | % Disease | N | % NVT | N | [illegible] |
| Gauteng | 2000-2014 | 3161 | 1559 | 1602 | 2424 | 76.7 | 149 | 994 | 31.45 | 251 | 16.10 | 743 | [illegible] |
| Western Cape | 2000-2014 | 880 | 532 | 348 | 880 | 100 | 85 | 184 | 20.91 | 76 | 14.29 | 108 | [illegible] |
| KwaZulu-Natal | 2000-2014 | 649 | 357 | 292 | 649 | 100 | 77 | 172 | 26.5 | 52 | 14.57 | 120 | [illegible] |
| Free State | 2000-2014 | 347 | 219 | 128 | 347 | 100 | 65 | 80 | 23.05 | 39 | 17.81 | 41 | [illegible] |
| Eastern Cape | 2000-2014 | 300 | 149 | 151 | 300 | 100 | 59 | 74 | 24.67 | 28 | 18.79 | 46 | [illegible] |
| Mpumalanga | 2000-2014 | 1308 | 128 | 1180 | 194 | 14.8 | 111 | 645 | 49.31 | 29 | 22.66 | 616 | [illegible] |
| North West | 2001-2014 | 116 | 66 | 50 | 116 | 100 | 40 | 23 | 19.8 | 8 | 12.12 | 15 | [illegible] |
| Northern Cape | 2001-2014 | 80 | 27 | 53 | 80 | 100 | 31 | 20 | 25 | 6 | 22.22 | 14 | [illegible] |
| Limpopo | 2001-2014 | 69 | 43 | 26 | 69 | 100 | 37 | 14 | 20.29 | 7 | 16.28 | 7 | [illegible] |
| Other Africa | 2000-2018 | 1157 | - | - | 754 | 65.2 | - | 262 | 22.6 | - | - | - | |
| Other Continents | 2000-2015 | 2944 | - | - | 1659 | 56.4 | - | 481 | 16.3 | - | - | - | |

**Table S2. Isolate breakdown by carriage and disease for the dominant GPSCs used in the phylogenies, the initial dataset used for the lineage level relative risk, dataset included in the fitness analysis and AMR for which in-silico AMR typing was performed.**

| Year | Dominant GPSCs | | Relative Risk Lineage Level | | Fitness Analysis & AMR | | |
|---|---|---|---|---|---|---|---|
| | Carriage | Disease | Carriage | Disease | Carriage | Disease | Unknown |
| 2000 | 0 | 68 | 0 | 186 | 0 | 186 | 0 |
| 2001 | 0 | 72 | 0 | 224 | 0 | 224 | 0 |
| 2002 | 0 | 75 | 0 | 221 | 0 | 221 | 0 |
| 2003 | 0 | 129 | 0 | 313 | 0 | 311 | 2 |
| 2004 | 0 | 166 | 0 | 417 | 0 | 417 | 0 |
| 2005 | 0 | 215 | 0 | 466 | 0 | 466 | 0 |
| 2006 | 0 | 190 | 0 | 419 | 0 | 417 | 2 |
| 2007 | 0 | 199 | 0 | 426 | 0 | 425 | 0 |
| 2008 | 0 | 186 | 0 | 408 | 0 | 407 | 0 |
| 2009 | 102 | 196 | 381 | 381 | 377 | 380 | 0 |
| 2010 | 134 | 136 | 365 | 311 | 365 | 310 | 1 |
| 2011 | 98 | 136 | 367 | 301 | 360 | 296 | 5 |
| 2012 | 73 | 189 | 349 | 425 | 349 | 325 | 7 |
| 2013 | 60 | 70 | 388 | 276 | 387 | 267 | 6 |
| 2014 | 0 | 81 | 0 | 286 | 0 | 285 | 1 |
| Totals | 467 | 2108 | 1850 | 5060 | 1838 | 4937 | 23 |
| Overall Totals | 2575 | | 6910 | | 6798 | | |

**Table S3.**

**Antimicrobial resistance across South Africa for three classes of antimicrobials. The antimicrobials investigated include** penicillin (beta-lactam), erythromycin, clindamycin (macrolides), and co-trimoxazole (sulfonamide). Some isolates may be resistant to more than one antibiotic.

| Antimicrobial | Resistant (%) | Resistant Carriage (%) | Resistant IPD (%) | Resistant VT (%) | Resistant (N) | NA |
|---|---|---|---|---|---|---|
| Penicillin_WGS | 48.18 | 44.12 | 49.59 | 90.35 | 3273 | 5 |
| Erythromycin_WGS | 17.3 | 15.63 | 17.86 | 93.36 | 1175 | 6 |
| Clindamycin_WGS | 11.15 | 7.63 | 12.41 | 95.90 | 757 | 6 |
| Cotrimoxazole_WGS | 68.39 | 69.76 | 67.82 | 81.50 | 4596 | 78 |
| Total | | | | | 6798 | |

**Table S4. GPSCs included in time-resolved phylogenetic trees including the total number and the number from South Africa for each.**

| Lineage | N (overall) | N (SA) |
|---|---|---|
| GPSC2 | 1394 | 507 |
| GPSC17 | 531 | 465 |
| GPSC14 | 512 | 413 |
| GPSC13 | 598 | 308 |
| GPSC5 | 831 | 300 |
| GPSC10 | 709 | 236 |
| GPSC1 | 1911 | 199 |
| GPSC79 | 95 | 83 |
| GPSC68 | 95 | 64 |

**Table S5. Relative Risk of pairs being within each distance, stratified by relatedness, compared to distant pairs which are >1000km apart.**

| | Carriage & Disease | | | Disease | | | |
|---|---|---|---|---|---|---|---|
| **Distance** | **mean** | **lowerCI** | **upperCI** | **mean** | **lowerCI** | **upperCI** | **Relatedness** |
| **Within Province** | 2.431 | 2.024 | 3.170 | 2.591 | 2.146 | 3.108 | 0 - 5 |
| **<500** | 1.543 | 1.254 | 1.976 | 1.676 | 1.340 | 2.140 | 0 - 5 |
| **500-1000** | 1.223 | 1.025 | 1.501 | 1.352 | 1.063 | 1.734 | 0 - 5 |
| **Distant Pairs** | 1.006 | 0.999 | 1.013 | 1.007 | 0.998 | 1.016 | 0 - 5 |
| **Other Africa** | 0.100 | 0.100 | 0.100 | 0.100 | 0.100 | 0.100 | 0 - 5 |
| **Outside Africa** | 0.100 | 0.100 | 0.100 | 0.100 | 0.100 | 0.100 | 0 - 5 |
| **Within Province** | 2.593 | 2.261 | 3.027 | 2.589 | 2.201 | 3.282 | 5-10 |
| **<500** | 1.861 | 1.576 | 2.311 | 1.792 | 1.535 | 2.190 | 5-10 |
| **500-1000** | 1.510 | 1.186 | 1.873 | 1.499 | 1.192 | 1.797 | 5-10 |
| **Distant Pairs** | 1.002 | 0.998 | 1.005 | 1.001 | 0.998 | 1.005 | 5-10 |
| **Other Africa** | 0.110 | 0.100 | 0.154 | 0.117 | 0.100 | 0.165 | 5-10 |
| **Outside Africa** | 0.100 | 0.100 | 0.100 | 0.100 | 0.100 | 0.100 | 5-10 |
| **Within Province** | 2.080 | 1.864 | 2.300 | 2.034 | 1.803 | 2.321 | 10-20 |
| **<500** | 1.709 | 1.513 | 1.892 | 1.645 | 1.464 | 1.851 | 10-20 |
| **500-1000** | 1.413 | 1.235 | 1.603 | 1.400 | 1.245 | 1.638 | 10-20 |
| **Distant Pairs** | 1.003 | 1.001 | 1.005 | 1.003 | 1.000 | 1.006 | 10-20 |
| **Other Africa** | 0.134 | 0.100 | 0.159 | 0.129 | 0.100 | 0.168 | 10-20 |
| **Outside Africa** | 0.429 | 0.376 | 0.491 | 0.424 | 0.371 | 0.496 | 10-20 |
| **Within Province** | 1.344 | 1.152 | 1.512 | 1.328 | 1.178 | 1.530 | 20 - 200 |
| **<500** | 1.697 | 1.450 | 1.880 | 1.655 | 1.443 | 1.951 | 20 - 200 |
| **500-1000** | 0.932 | 0.812 | 1.068 | 0.936 | 0.807 | 1.107 | 20 - 200 |
| **Distant Pairs** | 1.000 | 0.998 | 1.004 | 1.000 | 0.998 | 1.003 | 20 - 200 |
| **Other Africa** | 0.100 | 0.100 | 0.112 | 0.100 | 0.100 | 0.116 | 20 - 200 |
| **Outside Africa** | 0.141 | 0.121 | 0.164 | 0.139 | 0.110 | 0.154 | 20 - 200 |
| **Within Province** | 1.268 | 1.230 | 1.308 | 1.387 | 1.336 | 1.430 | Same Lineage |
| **<500** | 1.010 | 0.980 | 1.042 | 1.229 | 1.166 | 1.286 | Same Lineage |
| **500-1000** | 1.085 | 1.025 | 1.122 | 1.139 | 1.093 | 1.198 | Same Lineage |
| **Distant Pairs** | 1.005 | 1.004 | 1.006 | 1.001 | 1.001 | 1.002 | Same Lineage |

*For the by-lineage comparison the denominator is pairs which are from different lineages. We repeated this analysis including only pairs isolated from disease.*

**Table S6. Relative Risk of similarity across different distances and for a rolling window of relatedness (tMRCA).**

| **Distance Range** | MRCA | | **Carriage&Disease** | | | **Disease** | | |
|---|---|---|---|---|---|---|---|---|
| | **range** | **median** | **RR** | **lowerCI** | **upperCI** | **RR** | **lowerCI** | **upperCI** |
| **Within Province** | 0 - 1 | 0.5 | 2.561 | 1.899 | 4.030 | 1.680 | 1.222 | 2.394 |
| **<500** | 0 - 1 | 0.5 | 0.691 | 0.428 | 1.119 | 0.915 | 0.672 | 1.291 |
| **500-1000** | 0 - 1 | 0.5 | 1.385 | 0.893 | 2.088 | 1.057 | 0.948 | 1.784 |
| **Distant Pairs** | 0 - 1 | 0.5 | 1.031 | 1.006 | 1.060 | 1.016 | 1.003 | 1.034 |
| **Other Africa** | 0 - 1 | 0.5 | 0.011 | 0.009 | 0.018 | 0.012 | 0.009 | 0.016 |
| **Outside Africa** | 0 - 1 | 0.5 | 0.007 | 0.005 | 0.010 | 0.010 | 0.008 | 0.014 |
| **Within Province** | 0 - 11 | 5.5 | 3.030 | 2.611 | 3.389 | 2.665 | 2.325 | 2.962 |
| **<500** | 0 - 11 | 5.5 | 1.799 | 1.572 | 2.049 | 1.683 | 1.629 | 1.963 |
| **500-1000** | 0 - 11 | 5.5 | 1.679 | 1.392 | 1.932 | 1.308 | 1.085 | 1.456 |
| **Distant Pairs** | 0 - 11 | 5.5 | 1.012 | 1.008 | 1.016 | 1.002 | 0.998 | 1.006 |
| **Other Africa** | 0 - 11 | 5.5 | 0.108 | 0.076 | 0.149 | 0.103 | 0.064 | 0.131 |
| **Outside Africa** | 0 - 11 | 5.5 | 0.013 | 0.009 | 0.019 | 0.014 | 0.009 | 0.018 |
| **Within Province** | 0 - 21 | 10.5 | 2.237 | 1.996 | 2.363 | 2.206 | 2.069 | 2.333 |
| **<500** | 0 - 21 | 10.5 | 1.519 | 1.341 | 1.624 | 1.648 | 1.531 | 1.805 |
| **500-1000** | 0 - 21 | 10.5 | 1.533 | 1.373 | 1.687 | 1.405 | 1.246 | 1.527 |
| **Distant Pairs** | 0 - 21 | 10.5 | 1.011 | 1.008 | 1.013 | 1.003 | 1.002 | 1.004 |
| **Other Africa** | 0 - 21 | 10.5 | 0.123 | 0.098 | 0.149 | 0.116 | 0.104 | 0.133 |
| **Outside Africa** | 0 - 21 | 10.5 | 0.286 | 0.259 | 0.320 | 0.420 | 0.344 | 0.462 |
| **Within Province** | 0 - 31 | 15.5 | 2.241 | 2.123 | 2.456 | 2.264 | 2.079 | 2.441 |
| **<500** | 0 - 31 | 15.5 | 1.591 | 1.455 | 1.708 | 1.746 | 1.637 | 1.906 |
| **500-1000** | 0 - 31 | 15.5 | 1.641 | 1.520 | 1.824 | 1.483 | 1.391 | 1.555 |
| **Distant Pairs** | 0 - 31 | 15.5 | 1.012 | 1.010 | 1.014 | 1.002 | 1.001 | 1.004 |
| **Other Africa** | 0 - 31 | 15.5 | 0.154 | 0.123 | 0.183 | 0.131 | 0.121 | 0.146 |
| **Outside Africa** | 0 - 31 | 15.5 | 0.644 | 0.594 | 0.718 | 0.703 | 0.615 | 0.746 |
| **Within Province** | 0 - 41 | 20.5 | 2.550 | 2.385 | 2.810 | 2.587 | 2.321 | 2.966 |
| **<500** | 0 - 41 | 20.5 | 1.833 | 1.664 | 2.031 | 1.974 | 1.845 | 2.223 |
| **500-1000** | 0 - 41 | 20.5 | 1.639 | 1.444 | 1.820 | 1.484 | 1.416 | 1.651 |
| **Distant Pairs** | 0 - 41 | 20.5 | 1.012 | 1.010 | 1.015 | 1.003 | 1.003 | 1.006 |
| **Other Africa** | 0 - 41 | 20.5 | 0.146 | 0.119 | 0.174 | 0.116 | 0.103 | 0.135 |
| **Outside Africa** | 0 - 41 | 20.5 | 0.564 | 0.494 | 0.623 | 0.595 | 0.533 | 0.659 |
| **Within Province** | 0-51 | 26 | 2.655 | 2.427 | 2.967 | 2.771 | 2.507 | 2.964 |
| **<500** | 0-51 | 26 | 1.853 | 1.702 | 2.002 | 2.088 | 1.926 | 2.309 |
| **500-1000** | 0-51 | 26 | 1.702 | 1.552 | 1.802 | 1.516 | 1.344 | 1.605 |
| **Distant Pairs** | 0-51 | 26 | 1.015 | 1.013 | 1.017 | 1.004 | 1.004 | 1.007 |
| **Other Africa** | 0-51 | 26 | 0.139 | 0.115 | 0.162 | 0.099 | 0.083 | 0.110 |
| **Outside Africa** | 0-51 | 26 | 0.560 | 0.509 | 0.633 | 0.498 | 0.451 | 0.583 |
| **Within Province** | 11-61 | 36 | 2.312 | 2.088 | 2.527 | 2.384 | 2.111 | 2.616 |

| | | | | | | | | |
|---|---|---|---|---|---|---|---|---|
| **<500** | 11-61 | 36 | 1.742 | 1.560 | 1.901 | 1.943 | 1.737 | 2.081 |
| **500-1000** | 11-61 | 36 | 1.504 | 1.406 | 1.757 | 1.364 | 1.236 | 1.459 |
| **Distant Pairs** | 11-61 | 36 | 1.015 | 1.012 | 1.017 | 1.005 | 1.004 | 1.006 |
| **Other Africa** | 11-61 | 36 | 0.145 | 0.122 | 0.169 | 0.104 | 0.084 | 0.117 |
| **Outside Africa** | 11-61 | 36 | 0.812 | 0.735 | 0.877 | 0.480 | 0.448 | 0.580 |
| **Within Province** | 21 - 71 | 46 | 1.509 | 1.333 | 1.641 | 1.417 | 1.264 | 1.548 |
| **<500** | 21 - 71 | 46 | 1.224 | 1.120 | 1.311 | 1.154 | 1.097 | 1.268 |
| **500-1000** | 21 - 71 | 46 | 1.084 | 0.935 | 1.182 | 0.846 | 0.765 | 0.940 |
| **Distant Pairs** | 21 - 71 | 46 | 1.015 | 1.013 | 1.019 | 1.004 | 1.003 | 1.007 |
| **Other Africa** | 21 - 71 | 46 | 0.159 | 0.140 | 0.195 | 0.085 | 0.077 | 0.121 |
| **Outside Africa** | 21 - 71 | 46 | 0.589 | 0.541 | 0.646 | 0.272 | 0.233 | 0.316 |
| **Within Province** | 31 - 81 | 56 | 1.067 | 0.946 | 1.213 | 0.989 | 0.891 | 1.052 |
| **<500** | 31 - 81 | 56 | 0.859 | 0.760 | 0.963 | 0.808 | 0.772 | 0.875 |
| **500-1000** | 31 - 81 | 56 | 0.787 | 0.681 | 0.866 | 0.635 | 0.573 | 0.684 |
| **Distant Pairs** | 31 - 81 | 56 | 1.016 | 1.013 | 1.021 | 1.003 | 0.999 | 1.005 |
| **Other Africa** | 31 - 81 | 56 | 0.126 | 0.106 | 0.148 | 0.065 | 0.047 | 0.084 |
| **Outside Africa** | 31 - 81 | 56 | 0.376 | 0.332 | 0.421 | 0.107 | 0.095 | 0.120 |
| **Within Province** | 41 - 91 | 66 | 0.644 | 0.577 | 0.724 | 0.699 | 0.606 | 0.741 |
| **<500** | 41 - 91 | 66 | 0.724 | 0.638 | 0.825 | 0.647 | 0.568 | 0.701 |
| **500-1000** | 41 - 91 | 66 | 0.713 | 0.578 | 0.842 | 0.651 | 0.571 | 0.671 |
| **Distant Pairs** | 41 - 91 | 66 | 1.005 | 1.001 | 1.008 | 1.000 | 0.999 | 1.002 |
| **Other Africa** | 41 - 91 | 66 | 0.107 | 0.084 | 0.130 | 0.065 | 0.058 | 0.070 |
| **Outside Africa** | 41 - 91 | 66 | 0.270 | 0.226 | 0.320 | 0.095 | 0.083 | 0.127 |
| **Within Province** | 51 - 101 | 76 | 0.548 | 0.482 | 0.619 | 0.474 | 0.438 | 0.533 |
| **<500** | 51 - 101 | 76 | 0.680 | 0.594 | 0.766 | 0.461 | 0.403 | 0.521 |
| **500-1000** | 51 - 101 | 76 | 0.680 | 0.553 | 0.816 | 0.608 | 0.542 | 0.737 |
| **Distant Pairs** | 51 - 101 | 76 | 1.002 | 0.999 | 1.006 | 1.000 | 1.000 | 1.003 |
| **Other Africa** | 51 - 101 | 76 | 0.108 | 0.082 | 0.132 | 0.076 | 0.060 | 0.091 |
| **Outside Africa** | 51 - 101 | 76 | 0.271 | 0.229 | 0.316 | 0.113 | 0.109 | 0.123 |
| **Within Province** | 61 - 111 | 86 | 0.546 | 0.477 | 0.635 | 0.508 | 0.464 | 0.603 |
| **<500** | 61 - 111 | 86 | 0.678 | 0.552 | 0.762 | 0.501 | 0.433 | 0.581 |
| **500-1000** | 61 - 111 | 86 | 0.725 | 0.605 | 0.835 | 0.690 | 0.589 | 0.760 |
| **Distant Pairs** | 61 - 111 | 86 | 1.001 | 0.998 | 1.005 | 0.999 | 0.996 | 1.001 |
| **Other Africa** | 61 - 111 | 86 | 0.205 | 0.152 | 0.251 | 0.126 | 0.106 | 0.168 |
| **Outside Africa** | 61 - 111 | 86 | 0.319 | 0.264 | 0.371 | 0.189 | 0.166 | 0.213 |
| **Within Province** | 71 - 121 | 96 | 0.635 | 0.562 | 0.718 | 0.565 | 0.462 | 0.617 |
| **<500** | 71 - 121 | 96 | 0.780 | 0.682 | 0.894 | 0.577 | 0.497 | 0.637 |
| **500-1000** | 71 - 121 | 96 | 0.805 | 0.659 | 0.941 | 0.849 | 0.744 | 0.882 |
| **Distant Pairs** | 71 - 121 | 96 | 0.995 | 0.991 | 1.000 | 0.995 | 0.993 | 0.997 |
| **Other Africa** | 71 - 121 | 96 | 0.223 | 0.180 | 0.268 | 0.172 | 0.123 | 0.194 |

| | | | | | | | | |
|---|---|---|---|---|---|---|---|---|
| **Outside Africa** | 71 - 121 | 96 | 0.410 | 0.364 | 0.492 | 0.292 | 0.252 | 0.328 |

**Table S7. The relative risk of transmission chains being in each municipality after 1 year of transmission for all municipalities with a relative risk >1.**

| Municipality | Province | RR at 1 year |
|---|---|---|
| City of Johannesburg | Gauteng | 36.27234 |
| Ethekwini | KwaZulu-Natal | 27.36162 |
| City of Cape Town | Western Cape | 25.43112 |
| Ekurhuleni | Gauteng | 18.83115 |
| City of Tshwane | Gauteng | 17.84718 |
| Nelson Mandela Bay | Eastern Cape | 6.80979 |
| Buffalo City | Eastern Cape | 5.51343 |
| Mangaung | Free State | 4.67025 |
| Polokwane | Limpopo | 4.57314 |
| Thulamela | Limpopo | 3.72567 |
| Mbombela | Mpumalanga | 3.54978 |
| Rustenburg | North West | 2.92734 |
| Bushbuckridge | Mpumalanga | 2.67462 |
| Makhado | Limpopo | 2.67033 |
| King Sabata Dalindyebo | Eastern Cape | 2.64966 |
| City of Matlosana | North West | 2.27253 |
| Emalahleni.mp | Mpumalanga | 2.21247 |
| Greater Tzaneen | Limpopo | 2.08533 |
| Newcastle | KwaZulu-Natal | 2.00226 |
| Greater Tubatse | Limpopo | 1.93674 |
| uMhlathuze | KwaZulu-Natal | 1.76397 |
| Matjhabeng | Free State | 1.75656 |
| The Msunduzi | KwaZulu-Natal | 1.70235 |
| Nkomazi | Mpumalanga | 1.62864 |
| Mafikeng | North West | 1.54947 |
| Govan Mbeki | Mpumalanga | 1.27608 |
| Ngquza Hill | Eastern Cape | 1.21797 |
| Mogalakwena | Limpopo | 1.19574 |
| Emfuleni | Gauteng | 1.19145 |
| Nyandeni | Eastern Cape | 1.16298 |
| Makhuduthamaga | Limpopo | 1.15557 |
| Steve Tshwete | Mpumalanga | 1.08615 |
| George | Western Cape | 1.06938 |

**Table S8. Absolute fitness effects adjusted by proportion in each group**

| | prePCV | | | postPCV | | |
|---|---|---|---|---|---|---|
| Serotype Group | mean | lowerCI | upperCI | mean | lowerCI | upperCI |
| NVT | 1.06 | 1.02 | 1.1 | 1.31 | 1.21 | 1.43 |
| PCV7 | 0.99 | 0.97 | 1.03 | 0.8 | 0.73 | 0.88 |
| PCV13 | 0.99 | 0.95 | 1.03 | 0.9 | 0.81 | 0.99 |

**Table S9. Absolute fitness effects stratified by group for the model incorporating both vaccine status and antimicrobial resistance profile.**

| | pre-PCV | | | post-PCV | | | Denominator |
|---|---|---|---|---|---|---|---|
| Group | mean | lowerCI | upperCI | mean | lowerCI | upperCI | |
| VT_R | 1 | 1 | 1 | 0.99 | 0.97 | 1.01 | all VT |
| VT_S | 1 | 1 | 1 | 1 | 1 | 1.01 | all VT |
| NVT_R | 0.99 | 0.97 | 1 | 1.1 | 1.05 | 1.15 | all NVT |
| NVT_S | 1 | 1 | 1.01 | 0.98 | 0.96 | 1 | all NVT |
| VT_overall | 0.99 | 0.98 | 1 | 0.89 | 0.84 | 0.94 | all serotypes |
| NVT_overall | 1.05 | 1.02 | 1.07 | 1.16 | 1.09 | 1.24 | all serotypes |
| nvt_s | 1.13 | 1.06 | 1.22 | 1.09 | 1.01 | 1.18 | all serotypes and amr |
| nvt_r | 1.03 | 1 | 1.06 | 1.41 | 1.33 | 1.5 | all serotypes and amr |
| vt_s | 0.98 | 0.96 | 1 | 0.92 | 0.85 | 0.98 | all serotypes and amr |
| vt_r | 0.98 | 0.96 | 0.99 | 0.89 | 0.84 | 0.94 | all serotypes and amr |
